# The filopodial scaffold polyphosphate dictates cell adhesion-*versus*-invasion decisions

**DOI:** 10.64898/2026.09.01.747915

**Authors:** Akash Rai, Aanchal Jain, Hannah Zunker, Bonje N. Obua, Harshita Ramchandani, Jian Guan, Mousumi Akter, Yongkang Xi, Amanda Erwin, Pavithra Mahadevan, Bryndon J. Oleson, Adrian B.J. Bardwell, Jaclyn Shoudis, Jiage Song, Megha Suresh, Shyamal Mosalaganti, Allen P. Liu, Junior West, Ursula Jakob

## Abstract

Inorganic polyphosphate (polyP) is an ancient polymer conserved across all life, serving cell type- and location–specific functions in every major compartment. Yet its role at the plasma membrane, where it accumulates to peak levels in many primary cells, is largely unknown. Here we identify polyP as a stabilizing component of filopodia, actin-based membrane protrusions that govern cell adhesion, contact inhibition, and chemotaxis. Elevating cellular polyP increases filopodial stability and enhances cell adhesion, whereas reducing polyP accelerates filopodial disassembly and promotes cell migration. Mechanistically, we find that polyP acts as a structural filopodial scaffold, recruiting and organizing IRSp53, a membrane curvature–inducing protein. We show that metastatic fibroblasts and breast cancer organoids carry markedly reduced and intracellularly reorganized polyP levels relative to their non-transformed counterparts. Restoring endogenous polyP via lipid-nanoparticle delivery suppresses their invasive phenotypes and reverses pro-metastatic gene expression signatures, implicating polyP as a primordial tumor suppressor.

## Introduction

Inorganic polyphosphate (polyP) has long been regarded as one of Earth’s earliest energy-rich compound^1^. Comprised of chains with as few as three and as many as thousands of phosphoanhydride-linked phosphates, polyP is abundantly and universally present in all studied cells and organisms ^2,3^. Given its prebiotic origins, it is reasonable to infer that biological processes evolved in the context of this highly negatively charged polyanion. Indeed, over the past few decades, a growing body of literature has revealed a highly diverse functional repertoire of polyP in both prokaryotic and eukaryotic cells ^4^, with many of polyP’s functions specific to cell types, subcellular localization, and/or chain length, further contributing to its versatility ^5^. A notable example is the long-chain polyP found in platelets, which, upon its release, enhances blood clotting by influencing specific enzymes in the coagulation cascade ^6,7^. This function is exclusively attributed to polyP present in the dense granules of these cells ^8^. An early study examining the distribution of polyP in mammalian cells found its presence in the nucleus, cytosol, mitochondria, and along the plasma membrane ^9^. However, functional studies aimed at elucidating the roles of polyP in these various subcellular locations have lagged behind this initial discovery. This delay is primarily due to the challenges in manipulating endogenous polyP levels in mammalian cells. Recent evidence suggests that the F_0_F_1_-ATP synthase may be involved in mitochondrial polyP synthesis ^10^. However, inhibiting this machinery not only reduces polyP levels but also induces system-wide changes in ATP concentrations. Conversely, the known mammalian polyP-degrading enzymes hydrolyze the phosphoanhydride bonds of many different substrates ^11,12^, making it difficult to selectively reduce polyP levels without impacting other phosphorylated metabolites. As a result, disentangling the specific functional contributions of polyP amidst these systemic changes has been challenging. To address these limitations, two major strategies have emerged in recent years: i) the cytosolic overexpression of *Escherichia coli* polyphosphate kinase (EcPPK), which significantly increases cellular polyP levels without altering ATP concentrations ^13,14^, and ii) the overexpression of *Saccharomyces cerevisiae* exopolyphosphatase (ScPPX), which markedly decreases polyP levels in the cell compartments in which the enzyme is expressed ^15,16^. By utilizing these strategies, researchers have started to document the effects of altered polyP levels on transcriptional, proteomic, and physiological levels. These studies revealed that nuclear polyP plays a role in rRNA biogenesis through its interaction with RNA polymerase I ^17^ and regulates the fluidity of the nucleolus ^18^.

Mitochondrial polyP has been shown to influence cellular ATP homeostasis, calcium signaling, and ion channel formation ^19–21^ while much of the cytosolic polyP is stored in specialized granules and endolysosomal compartments, where it appears to be involved in phosphate homeostasis and pH buffering ^22^. In contrast to these studies, the role of polyP along the plasma membrane, which exhibits some of the highest polyP levels in primary cells ^9^, has remained largely enigmatic. It has been shown that exogenous polyP supplementation enhances epithelial barrier function via a mechanism that is mediated by integrin-β1 and increased p38 phosphorylation and accompanied by enhanced stability of F-actin and E-cadherin ^23^ and promotes increased stress fiber formation ^24^. These findings suggested that polyP may play a role in modulating cellular actin dynamics and/or cell adhesion.

Here we demonstrate that genetically elevating or depleting polyP regulates the number, length and dynamic stability of filopodia, which are actin-based cell protrusions involved in cell adhesion and migration. We find that polyP interacts with the Inverse Bin-Amphiphysin-Rvs (IBAR)-domain protein IRSp53, whose assembly into higher-order oligomeric states induces membrane curvature, a critical step in filopodia formation. Functionally, polyP levels correlate positively with cell adhesion and inversely with cell migration. Metastatic fibroblasts and breast cancer organoids show reduced polyP levels, and lipid-nanoparticle delivery restores filopodia, suppresses invasion, and reverses pro-metastatic signatures, establishing polyP as a membrane scaffold with anti-invasive potential.

## Results

### Cellular polyP levels positively correlate with filopodia number, length, and stability

To investigate the functional implications of polyP’s localization at the plasma membrane, we examined NIH3T3 mouse fibroblasts, the original mammalian cell line used to characterize intracellular polyP distribution^9^, alongside human BJ fibroblasts for comparison. We visualized polyP upon fixation using the fluorescently tagged polyP-binding probe PPXBD-GFP^25^. Consistent with previous reports, both cell lines showed polyP broadly distributed throughout the cytosol (Fig. 1A, B; S1A, B). In addition, however, we noticed distinct polyP puncta along the plasma membrane and, even more strikingly, along the longitudinal axis of finger-like structures projecting from the cell surface into the surrounding environment (*insets*, Fig. 1A, B). These actin-positive protrusions resembled filopodia, well-conserved structures known to mediate cell adhesion, chemotaxis, and contact inhibition^26–29^. Staining both cell lines for the non-canonical motor protein myosin-X (MyoX), a driver of filopodia formation that localizes to their tips^30^, revealed that the ends of each polyP-positive protrusion were MyoX–positive (Fig. 1A, B; S1A, B). These findings provided the first evidence that polyP might constitute a previously unrecognized component of mammalian filopodia or filopodia-like structures. To investigate whether and how polyP influences the morphology, formation, and/or stability of these protrusions, we transfected NIH3T3 cells with the *E. coli* polyP-kinase EcPPK1. We confirmed a significant increase in cellular polyP levels compared to mock-transfected control cells by both image-based (Fig. 1C, D; S1B) and biochemical quantification (Fig. 1E). Moreover, we observed that EcPPK1-transfected NIH3T3 cells exhibit a significant increase in both the number (Fig. 1F) and average length (Fig. 1G) of protrusions that stain positively for polyP and MyoX relative to mock-transfected cells. Western blot analysis of these cell lysates using antibodies against MyoX, CDC42 or IRSp53, each of which previously shown to increase filopodia formation when overexpressed^31–33^, revealed no significant difference in their steady-state levels following EcPPK1 expression (Fig. S1C-E). Moreover, and fully consistent with previous reports^14^, we did not observe any significant differences in cellular ATP (the proposed substrate of the polyP synthesis machinery)^10^ between mock- and EcPPK1-transfected NIH3T3 cells (Fig. S1F). These results excluded changes in the cellular energy state as a possible cause for the observed differences in filopodia formation. To independently validate that increased polyP levels increase the number and/or length of filopodia in mammalian cells, we tested human bone osteosarcoma epithelial U2OS cells, whose transfection efficiency is significantly higher than that of primary BJ cells. Similar to NIH3T3 cells, expression of EcPPK1 led to a significant polyP accumulation (Fig. S1G, H) and a pronounced increase in MyoX-positive filopodia-like structures (Fig. S1G, I). We also explored the effects of increased polyP levels in COS-7 fibroblasts, which form primarily lamellipodia under standard cultivation conditions, making them a suitable model for identifying novel inducers of filopodia formation^30,34^. Upon increasing the cellular polyP levels through the expression of EcPPK1 (Fig. 1H-J), we observed a dramatic alteration in the plasma membrane topology. While mock-transfected COS-7 cells displayed the characteristic smooth and ruffled surfaces with few detectable filopodia (Fig. 1H, upper panel), EcPPK1-expressing cells exhibited numerous MyoX and polyP-positive protrusions (Fig. 1H, lower panel). These results confirmed that COS-7 cells have the machinery to form filopodia and revealed that an increase in endogenous polyP levels is sufficient to trigger their formation. To finally test whether the observed effects of polyP on the number and/or length of filopodia in NIH3T3 cells might be due to polyP-mediated changes in the dynamics by which filopodia assemble and disassemble as they migrate along their substrate ^26,35^, we stably transfected the cells with the fluorescent actin probe LifeAct-iRFP670 ^36^ and conducted live-cell imaging. We found that in contrast to mock-transfected NIH3T3-LifeAct-iRFP670 cells, in which filopodia continuously form and retract, filopodia in cells expressing EcPPK1 show little to no apparent change in overall length over at least 5 min of measurements (Fig. 1L, Movie S1). Subsequent treatment of the cells with a low dose (250 nM) of latrunculin B, an inhibitor that prevents the addition of monomeric G-actin to the growing end of F-actin filaments^37,38^, confirmed this polyP-mediated increase in filopodial stability. We found that the filopodial F-actin dissociation rate in EcPPK1-expressing cells is at least three times slower (from 3.5 min to > 11 min) than in mock-transfected cells (Fig. 1M, N). These experiments, which revealed that filopodia formed in cells with higher-than-normal polyP concentrations are drastically more stable, provided a suitable explanation for why these cells harbor more and longer filopodia than cells with unaltered polyP levels.

**Figure 1.**
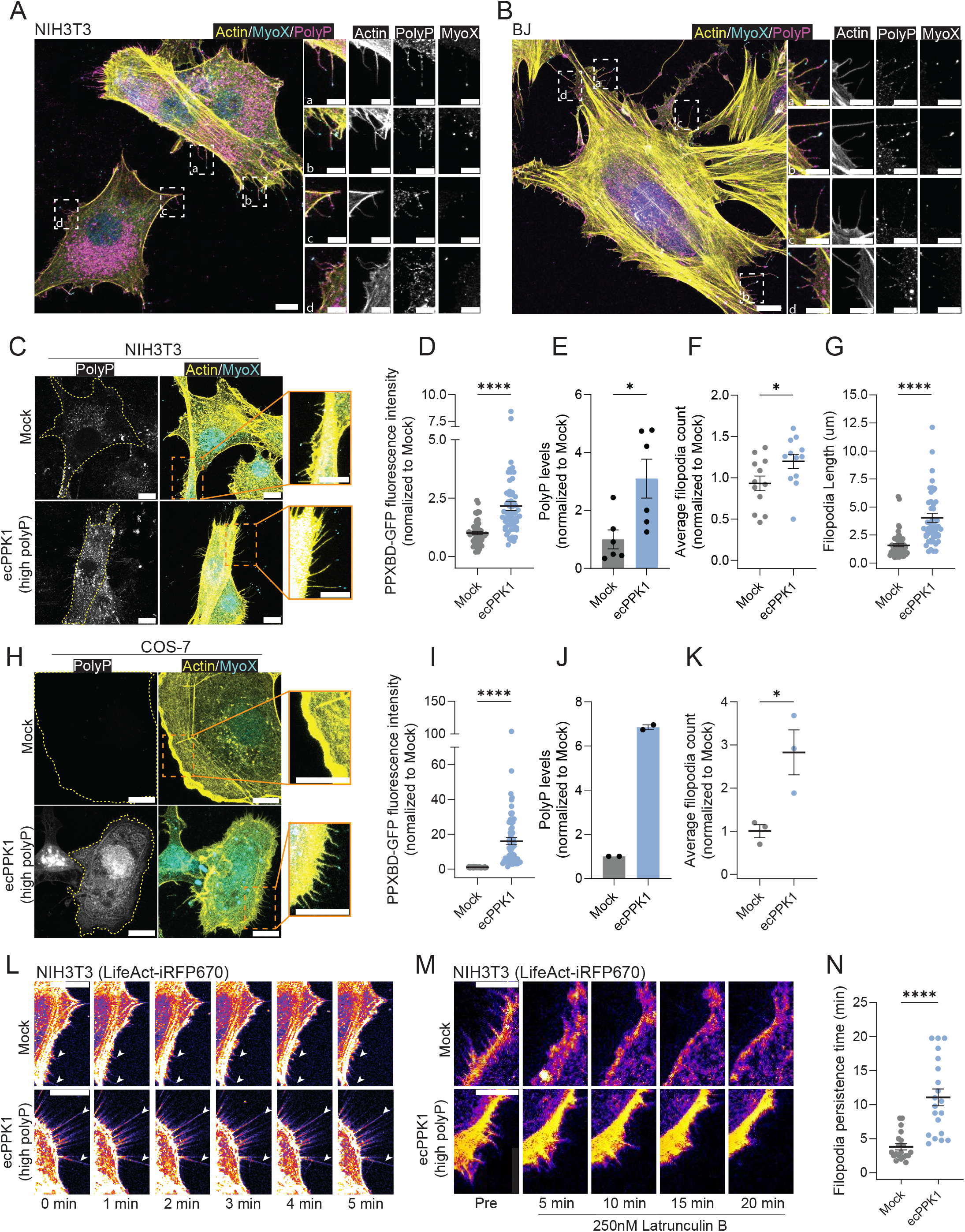
Increase in cellular polyP increases filopodia number, length and dynamics. Fluorescent images of (**A**) NIH3T3 mouse embryonic fibroblasts, (**B**) BJ human fibroblasts or **C**) (**C**) mock- or EcPPK1-transfected NIH3T3 fibroblasts were fixed and stained for polyP (magenta), actin (yellow) of MyoX (cyan). Individual regions of the membrane are shown enlarged. Quantification of the (**D**) fluorescent polyP signal, (**E**) extracted polyP, (**F**) filopodia number or (**G**) filopodia length of cells shown in (**C**). (**H**) Mock- or EcPPK1-transfected COS-7 fibroblasts were fixed and stained for polyP (gray), actin (yellow) or MyoX (cyan). Quantification of (**I**) fluorescent polyP signal, (**J**) extracted polyP or **(K**) filopodia number of cells shown in (**H**). (**L**) To observe filopodia formation and retraction, mock- or EcPPK1-transfected NIH3T3-LifeAct-iRF670 fibroblasts were live-imaged over 5 min. Still images of movie S1 are shown. White arrowheads depict individual filopodia. (**M**) Mock- or EcPPK1-transfected NIH3T3-LifeAct-iRF670 fibroblasts were live-imaged for 10 min (pre) before addition of 250 nM Latrunculin-B to assess filopodial retraction speeds. Still images of movie S2 are shown. (**N**) Quantification of filopodial retraction speed using filopodia of similar lengths of cells shown in (**M**). Scale bars: 10 µm (main), 5 µm (insets). Datapoints in panels **D, I**: PPXBD-GFP fluorescence intensity of individual cells (n=15-20 cells per replicate, 3 replicates) with mean fluorescence intensity of mock-transfected cells set to 1. Datapoints in panels **E, J**: biological replicates (n=3) of ∼200,000 cells used for polyP extraction and normalized to total protein content; Datapoints in panels **F, K:** average MyoX-positive filopodia count per cell (n=5-25). Datapoints in panel **N**: Retraction speed of individual filopodium tracked over time. A maximum of 5 filopodia per cell (n=3-4 cells) was used. Unpaired t-test was used to determine statistical significance between samples. * *p* < 0.05, *** *p* < 0.001, **** *p* < 0.0001. The data represented in panels D, F, G, I, K, N came from blinded experiments.

### Decrease in endogenous polyP levels increases the dynamic properties of filopodia

To investigate whether the reverse relationship is also correct, *i.e.*, whether decreasing the cellular levels of polyP decreases filopodia length and increases filopodial dynamics, we generated monoclonal NIH3T3 cell lines expressing ScPPX. To regulate the *in vivo* polyphosphatase activity of ScPPX, we fused the flag-tagged version of the enzyme to a destabilizing domain (DD) degron tag, generating the DD-3xFLAG-scPPX construct. Analysis of three individual monoclonal cell lines (Fig. S2A) confirmed that ScPPX is continuously degraded in the absence of its stabilizer Shield (Fig. S2B). Addition of Shield to the media, however, led to the accumulation of the ScPPX protein (Fig. S2B, C) and a significant decrease in intracellular polyP levels within 24 hours (Fig. 2A-C; S2A). Analysis of the membrane morphology of ScPPX-expressing NIH3T3 cells revealed that reducing the cellular polyP levels causes an approximately 40% reduction in the average filopodia number (Fig. 2D) and length (Fig. 2E) compared to mock-transfected cells. Again, and fully consistent with published reports^14^, we did not observe any significant differences in cellular ATP levels between ScPPX-transfected NIH3T3 cells grown with or without Shield (Fig. S1D), excluding the possibility that the observed changes are due to variations in the cellular energy state. To test whether these phenotypic changes reflect increased rates of filopodia formation and retraction, we stably transfected LifeAct-iRFP670 expressing NIH3T3 fibroblasts with the DD-3xFLAG-ScPPX construct. Live-cell imaging of polyP-depleted cells (i.e., cells cultivated with Shield) revealed a significant reduction in filopodial persistence times compared to similarly sized filopodia in cells grown without Shield (Fig. 2F, G; Movie S2). Based on these results, we concluded that the cellular concentration of polyP directly impacts the number, length, and dynamic properties of filopodia in fibroblasts.

**Figure 2.**
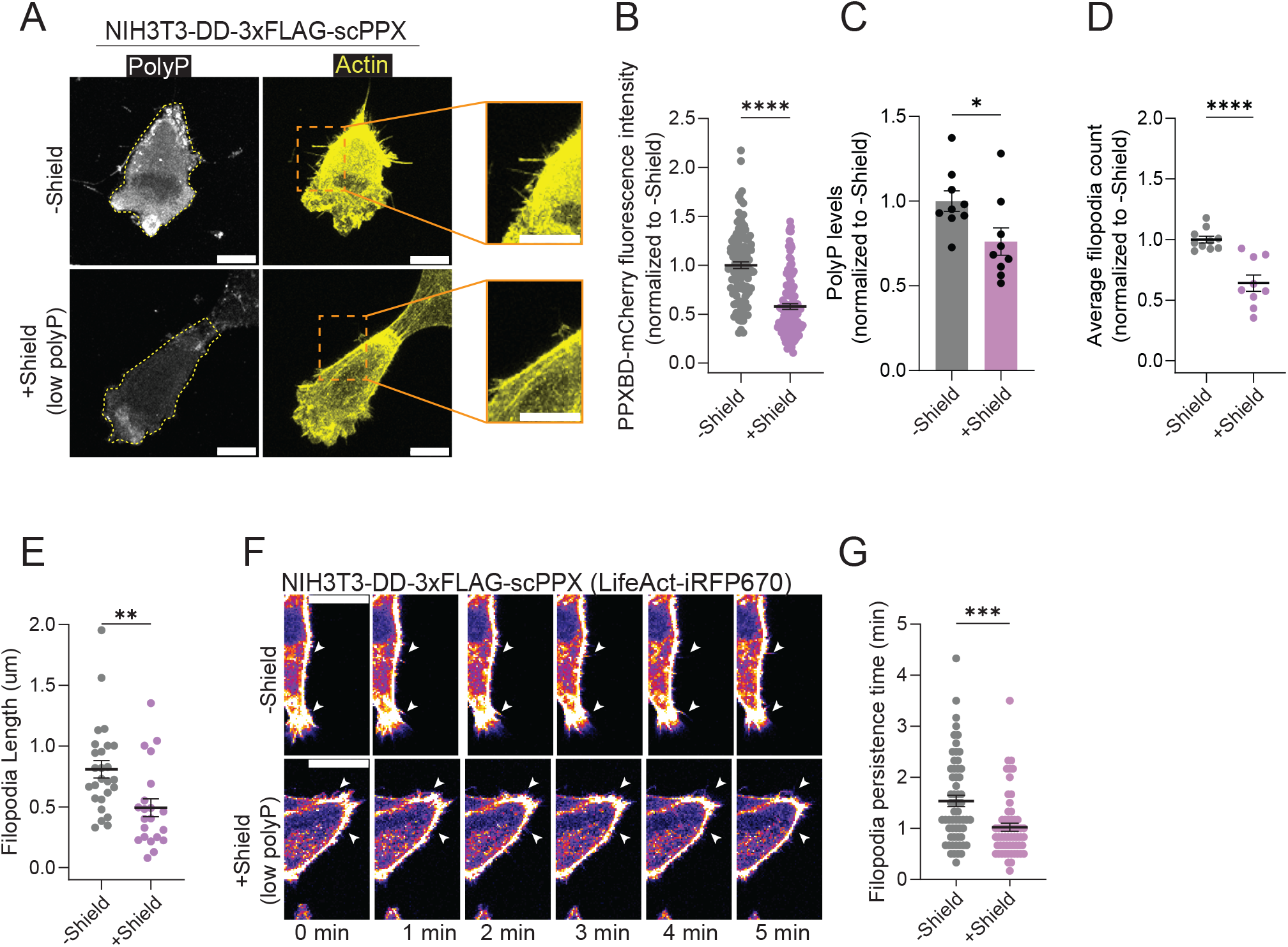
Decrease in cellular polyP levels increases filopodia dynamics. (**A**) NIH3T3 fibroblasts stably transfected with a 3xFLAG and degron (DD)-tagged yeast exopolyphosphatase (ScPPX) (DD-3xFLAG-ScPPX) and incubated in the absence or presence of the stabilizer Shield for 24h were fixed and stained for polyP (gray), actin (yellow) of MyoX (cyan). Quantification of the (**B**) fluorescent polyP signal, (**C**) extracted polyP, (**D**) filopodia number or (**E**) filopodia length of cells shown in (**A**). (**F**) NIH3T3-DD-3xFLAG-ScPPX fibroblasts expressing LifeAct-iRFP670 and cultivated in the absence (normal polyP levels) or presence of Shield (low polyP levels) were live-imaged. Still images of movie S3 are shown. White arrowheads depict individual filopodia. (**G**) Quantification of persistence times using filopodia of similar lengths of cells shown in (**F**). Scale bars for main images are 10 µm; inset scale bars are 5 µm. For information on individual data points shown in panels B, C, D, E and G please see legends to Figure 1D, E, F, G and N, respectively. Unpaired t-tests were used to determine statistical significance between samples. * *p* < 0.05, *** *p* < 0.001, **** *p* < 0.0001. The data represented in panels B,D,E,G came from blinded experiments.

### PolyP colocalizes and interacts with select filopodial proteins *in vivo*

Cryo-electron tomography (cryo-ET) analysis of individual filopodia in mock- and EcPPK1-expressing NIH3T3 cells revealed the presence of a visibly denser protein network between the filopodial membrane (Fig. 3A, yellow arrows) and the central actin filaments (Fig. 3A, blue arrows) in cells with higher-than-control polyP levels. Quantitative analysis of these membrane-associated densities (Fig. 3A, red arrows) showed that these differences are significant (Fig. 3B). These results, together with our finding that the observed changes in filopodia number and dynamics are not accompanied by an upregulation of key filopodial proteins suggested that polyP might act as a scaffolding agent, supporting the assembly of one or more filopodia-associated proteins into higher oligomeric structures along the membrane. This conclusion aligned well with previous studies demonstrating that polyP interacts with and stabilizes oligomerization-competent proteins in high-molecular-weight (HMW) assemblies ^39^. Many filopodial proteins work in the form of HMW structures; for instance, filopodia formation is initiated by IRSp53, an I-BAR protein, whose oligomerization in phosphatidylinositol-5,4-bisphosphate (PIP2)-enriched membrane regions induces convex membrane curvature^40,41^.

**Figure 3.**
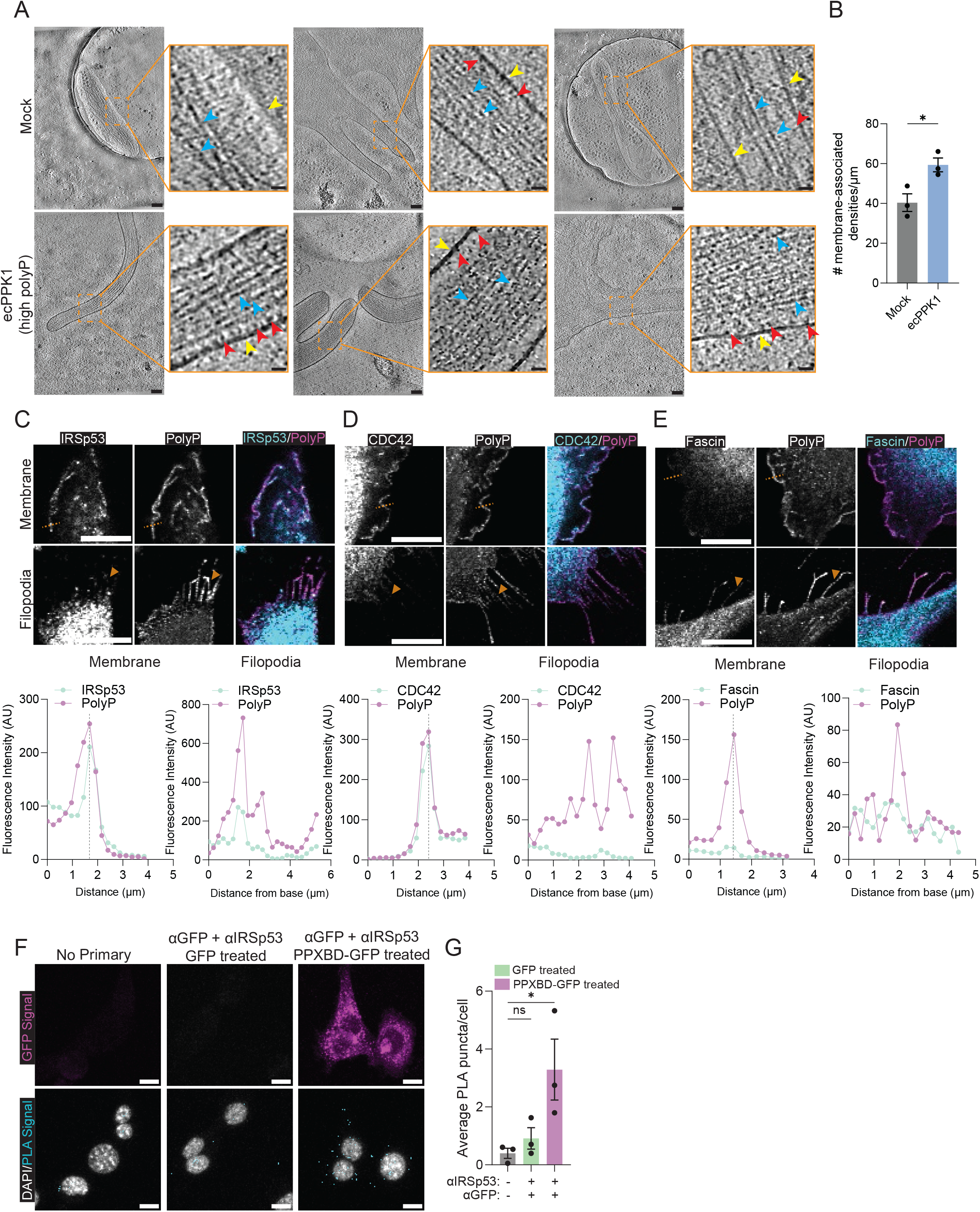
PolyP is a *hitherto* unknown component of mammalian filopodia. (**A**) Cryo-electron tomography of filopodia in mock (top)- or ecPPK1 (bottom)-transfected NIH3T3 fibroblasts. Three central slices through the reconstructed tomograms are shown in each case. Scale bars are 100 nm. Insets (orange boxes) depict zoomed-in views of the indicated filopodia regions; membrane (yellow arrows), actin bundles (blue arrows), and membrane-associated densities (red arrows). Scale bars for insets are 20 nm. (**B**) Quantification of the number of membrane-associated densities per µm in filopodia shown in (**A**). Each data point represents the total number of membrane densities counted along the length of a filopodium and divided by the total length; n=3 filopodia per cell (n=3 cells per condition). NIH3T3 fibroblasts were fixed and stained for polyP and either (**C**) IRSp53, (**D**) CDC42 or (**E**) fascin. False coloring was applied as shown above the panels. Yellow dotted lines indicate regions along the membrane or filopodia that were used for intensity profile analysis shown in lower panels. Additional examples can be found in Fig. S3A-C. (**F**) Proximity ligation assay (PLA) of fixed NIH3T3 fibroblasts. Cells were incubated with buffer alone (left), purified GFP (center) or purified PPXBD-GFP (right) followed by incubation with mouse anti-GFP and rabbit anti-IRSp53 antibodies. All samples were then incubated with the respective oligonucleotide-labeled secondary antibodies. The GFP signal is shown in magenta while the PLA amplification signal is depicted as cyan puncta. See Fig. S3I,J for controls. **(G**) Quantification of the number of positive puncta per cell per condition. Each data point represents the total number of PLA puncta in a field of view divided by the total number of cells in that same field of view (n>120 cells per condition per replicate; 3 replicates). Scale bars for all IF images are 10 µm. One-way ANOVA was used to determine statistical significance; ns, non-significant; * *p* < 0.05. Quantitative image analysis for panels B,G was conducted blinded.

Filopodia formation continues with the IRSp53-mediated recruitment of the small GTPase CDC42^40–42^ and the subsequent association with profilin, which drives rapid actin polymerization. The latter is further facilitated by MyoX, the motor protein that transports essential components to the growing tip ^30^. fascin stabilizes the 15-30 parallel actin bundles that form the core of the mature filopodia while IRSp53 tethers them to the membrane ^43^. To investigate whether polyP colocalizes with any of these key filopodial components, we co-stained fixed NIH3T3 cells with the polyP-binding protein PPXBD-GFP and antibodies against select filopodial proteins. We observed reproducible co-localization between polyP foci and IRSp53 (Fig. 3C, S3A), CDC42 (Fig. 3D, S3B) and fascin (Fig. 3E, S3C), and only sporadic colocalization between polyP foci and ARP2, MyoX or paxillin (Fig. S3D-F). Intensity profile analysis confirmed the colocalization between polyP and IRSp53 foci across the plasma membrane and along the filopodial axis (Fig. 3C, S3A) while colocalization between polyP and CDC42 was exclusively detected across the plasma membrane (Fig. 3D; S3B) and with fascin exclusively along the filopodial axis (Fig. 3E, S3C). These co-localizations were fully consistent with their respective functions in filopodia formation. However, we reasoned that polyP’s interactions with IRSp53 might be particularly relevant for its cellular effects as this protein plays a role in both the initial filopodia formation and in filopodia stability. Upon confirming that the colocalization between polyP and IRSp53 is not cell-type specific but also observed in BJ fibroblasts (Fig. S3D, E), we conducted a proximity ligation assay (PLA) to test whether polyP and IRSp53 come sufficiently close to interact. PLA is a highly specific and sensitive technique to detect and quantify protein-protein interactions at endogenous levels in fixed cells or tissues ^44^. As a positive control, we stained fixed NIH3T3 cells with oligonucleotide-labeled antibodies against IRSp53 and its known interaction partner actin, which yielded numerous amplification events (Fig. S3F, G). Importantly, we observed similar numbers of bright fluorescent puncta when we first incubated cells with PPXPD-GFP to stain for polyP, followed by incubation with oligonucleotide-linked anti-GFP and anti-IRSp53 antibodies (Fig. 3F, G). In contrast, we did not detect significant amplification events when we incubated the fixed cells with purified GFP instead of PPXPD-GFP and followed by incubation with the oligonucleotide-linked anti-GFP and anti-IRSp53 antibodies or when we added the secondary antibodies only (Fig. 3F, G). This interaction would position polyP at the PIP2-enriched membrane regions that are known to accumulate IRSp53 ^45^ and might explain polyP’s previously observed interaction with PIP4K2B, the kinase responsible for PIP2 generation in eukaryotic cells ^45^.

### PolyP promotes IRSp53 condensate formation *in vitro*

Central to the function of IRSp53 is its I-BAR domain, which mediates the association of IRSp53 into dimers at PIP2-enriched membranes ^40–42^. The assembly of several IRSp53 dimers into distinctly curved higher oligomeric states induces membrane curvature and precedes the recruitment of select regulators, such as CDC42 and the initiation of filopodia formation ^32,33,46,47^. Our finding that polyP increases filopodia formation without upregulating the levels of IRSp53 made us now wonder whether polyP serves as a scaffolding factor that promotes the clustering of existing IRSp53 molecules along the membrane. To test this idea, we purified IRSp53 and attempted to measure polyP-protein binding in solution. Unexpectedly, however, we found that a clear solution of 10 µM IRSp53 in 50 mM KPi, 100 mM KCl, pH 7.5 buffer turned immediately turbid upon addition of as little as 5 µM of the 300-mer polyP chain (all polyP concentrations given in Pi-units) (Fig. 4A). We saw very similar effects when we added the shorter 130-mer polyP and slightly reduced effects when we added the 60-mer instead. Of note, addition of the 14-mer polyP at the same total Pi-concentration did not cause any detectable change in turbidity, excluding the possibility of non-specific charge-effects. Consistent with earlier work which showed that polyP, in a chain length dependent manner, causes protein phase separation both *in vitro* and *in vivo*^39^ as well as published work that documented the droplet formation of purified IRSp53 in the presence of select binding partners such as PSD-95^48^, we now speculated that polyP might cause IRSp53 condensate formation. Analysis of select IRSp53-polyP samples by microscopy agreed with this conclusion and revealed that whereas IRSp53 alone failed to form any detectable droplets at concentrations up to 25 µM, presence of polyP-chains larger than 14 Pi-units caused significant droplet formation (Fig. 4B, Fig. S4A). A phase diagram of IRSp53 confirmed that the presence of polyP-300 reduces the c_sat_ of IRSp53 from over 25 µM to less than 500 nM (Fig. 4C). Mixing fluorescently labeled IRSp53-AF488 with fluorescently tagged polyP300-AF647 revealed that the two components form co-condensates and confirmed that neither component forms droplets on its own under these buffer conditions (Fig. 4D). The droplets were stable at physiological salt concentrations and only visibly dissolved between 100 and 200 mM NaCl (Fig. S4B). Fluorescence recovery after photobleaching (FRAP) experiments confirmed the fluidic nature of these droplets and, by comparing the fluidity of IRSp53 droplets formed with either the 130-mer or the 300-mer of polyP, suggested that the longer polyP chains further reduce the fluidity of the droplets (Fig. 4E). These findings led us to postulate that long-chain polyP assists in the formation and stabilization of filopodia through the scaffolding of multiple IRSp53 dimers along the membrane.

**Figure 4.**
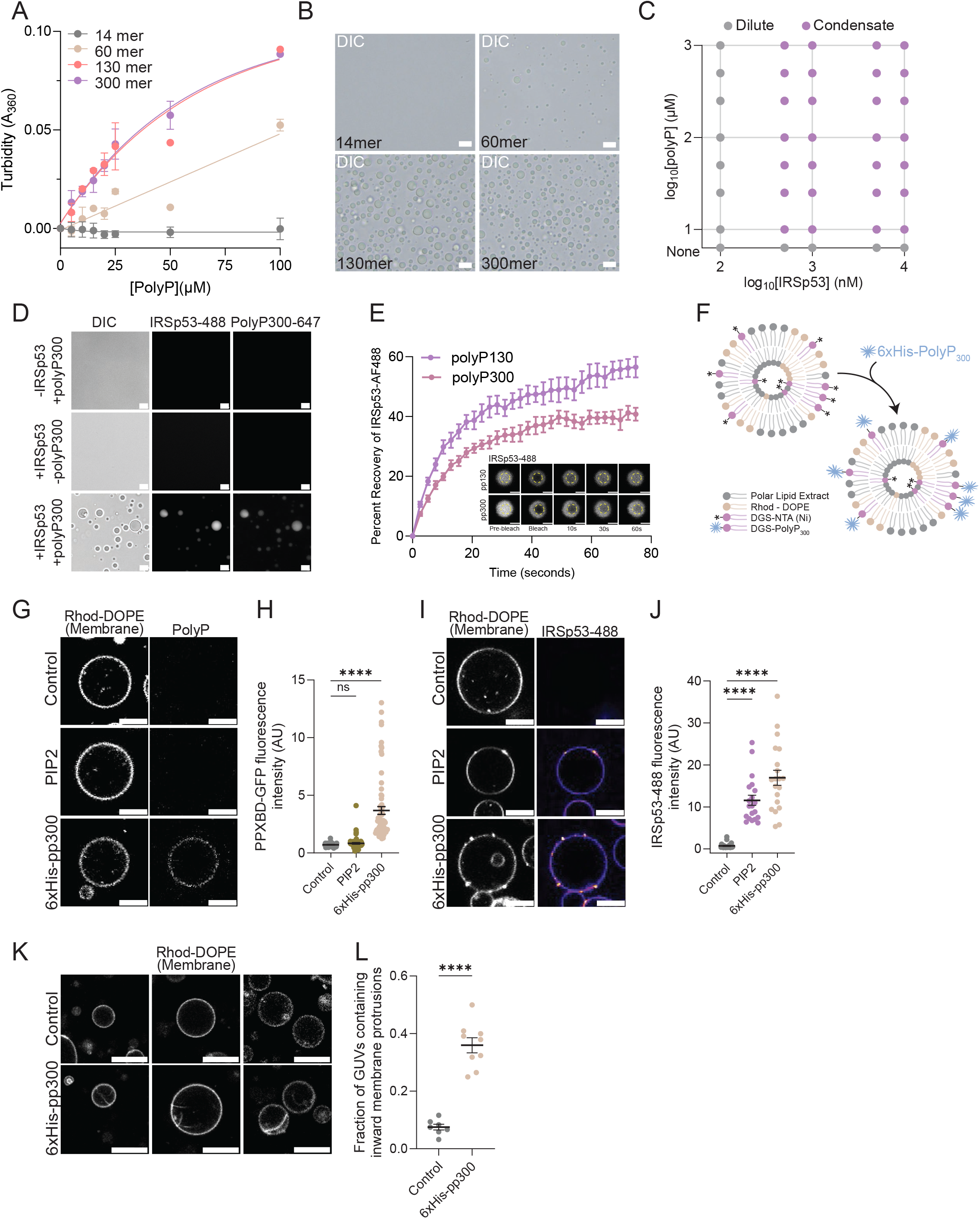
Membrane-bound polyP recruits IRSp53 and supports *ex vivo* filopodia formation. (**A**) Turbidity (OD360) of solutions containing 10 µM IRSp53 in 50 mM KPi, 100 mM KCl, pH 7.4 and increasing concentrations of select-length polyP chains (concentration given in Pi-units). (**B**). DIC images of 10 µM IRSp53 in the presence of 100 µM (in Pi-units) polyP-14, polyP-60, polyP-130 or polyP-300. Conditions as shown in (**A**). Scale bars: 10 µm. (**C**) Phase diagram of the indicated concentrations of IRSp53 and polyP-300. Conditions as shown in (**A**). Condensate formation was analyzed by microscopy. **(D**) Fluorescence microscopy images of solutions containing 10 µM IRSp53 (+ 2% AF488-IRSp53) and/or 100 µM polyP-300 (+ 2% AF647-polyP). Scale bars: 10 µm. (**E**) Fluorescence recovery upon photobleaching (FRAP) analysis of similarly sized droplets of 10 µM IRSp53 (+ 2% IRSp53-488) formed in the presence of either 100 µM (in Pi-units) polyP-130 or polyP-300. Inset: Representative images of FRAP experiment. Scale bars: 5 µm. (**F**) Illustration of the generation of polyP-enriched Giant Unilamellar Vesicles (GUVs). See materials and methods for details. (**G**) Membrane-associated polyP signal of control, PIP2-enriched or Ni-NTA enriched GUVS after incubation with 2.5 µM 6xHis-polyP-300. The membranes of GUVs were visualized using the rhodamine-DOPE signal while polyP was visualized upon staining with PPXBD-GFP. Scale bars: 10 µm. (**H**) Quantification of the polyP signal on the membranes of GUVs shown in (**G**). Each data point represents the PPXBD-GFP mean fluorescence intensity accumulating on the surface of a single GUV. n>30 GUVs were analyzed per condition per replicate. (**I**) IRSp53-488 signal on the membrane of control, PIP2-enriched or polyP-enriched GUVs after 15 min incubation with 50 nM IRSp53-488. The membrane signal is shown on the left. Scale bars: 10 µm. (**J**) Quantification of the IRSp53 signal on the membranes of GUVs shown in (**I**). Each data point represents the mean fluorescence intensity of IRSp53-488 on the surface of a single GUV. Results obtained with two separate GUV preparations are shown in Fig. S4H-K. (**K**) Fluorescence microscopy images of three different control or polyP-enriched GUV preparations upon incubation with IRSp53, VASP, profilin, fascin, capping protein, actin and ATP in filopodia forming-buffer for 15 mins. (**L**) Quantification of the number of inward filopodia of GUVs shown in (**K**). Each data point represents the percentage of GUVs with at least one single inward protrusion in a given field of view combining data from three independent GUV preparations. GUVs less than 5 µm in diameter were excluded from this analysis. Quantitative image analysis for panels H,J,L was conducted blinded. One-way ANOVA was used to determine statistical significance between samples in H, J. Unpaired t-tests were used to determine statistical significance between samples in L. ns, non significant; **** *p* < 0.0001

### PolyP-mediated IRSp53 recruitment to membranes promotes filopodia formation

To test whether the presence of long-chain polyP at the plasma membrane is sufficient to recruit IRSp53 and induce filopodia formation, we generated synthetic lipid bilayer structures in the form of giant unilamellar vesicles (GUVs). Consisting of a single lipid bilayer, whose composition can be precisely controlled, GUVs have previously been used to monitor filopodia formation *in vitro* ^42^. We mixed brain polar lipid extracts with rhodamine-red labeled dioleoyl phosphatidylethanol amine (DOPE) lipids to visualize the membrane. Next, we either added nothing (control), 1,2-dioleoyl-sn-glycero-3-phospho-L-serine, which generates the negative charge typically observed on the inner leaflet of membranes^49^ (PS), phosphatidylinositol 4,5-bisphosphate (PIP2), the phospholipid known to recruit IRSp53 to membranes^40,42^ or 18:1 DGS-NTA(Ni), a specialized nickel-chelating phospholipid for subsequent enrichment of His-tagged polyP-300^50^ (Fig. 4F). We conducted electroformation or inverted emulsion to generate GUVs ^51^, which are easily visible by the fluorescent rhodamine signal along the membrane (Fig. 4G, top panel; Fig. S4C). To generate polyP-enriched membranes, we next incubated DGS-NTA(Ni)-supplemented GUVs with 2.5 µM 6xHis-tag polyP-300^50^ (Fig. 4F). As controls, we also incubated control or PIP2-enriched GUVs with His-tagged polyP-300. Image analysis of the GUVs upon incubation with the polyP binding probe PPXBD-GFP revealed a pronounced enrichment of the GFP-signal only on the DGS-NTA(Ni)-supplemented GUVs incubated with His-tagged polyP but not on the other types of GUVs (Fig. 4G, H). These results gave us confidence that we could generate polyP-enriched synthetic membranes. To test whether these membranes recruit IRSp53, we next incubated the various GUVs with 50 nM fluorophore labelled IRSp53-488. We observed a significant increase in the recruitment of IRSp53-AF488 to GUVs enriched for polyP and PIP2 (Fig. 4I, J; Fig. S4C-F) but not for the control or PS-enriched GUVs (Fig. S4G,H), the latter fully consistent with published results^42^. Moreover, we also noted the previously observed clustering of IRSp53 on both PIP2-enriched^42^ as well as polyP-enriched membranes (Fig. 4I, S4C, E). Although the quantification of the number and size of IRSp53 clusters revealed that the ones formed on polyP-enriched membranes are on average not only of higher intensity (Fig. S4I) but also larger than those seen on PIP2-enriched membranes (Fig. S4J), we cannot exclude the possibility that these differences are due to relative differences in the levels or distribution of PIP2 and polyP on the artificial membranes. Nonetheless, these results demonstrated that polyP recruits and scaffolds IRSp53 clusters at the membrane. To finally test whether this recruitment is sufficient to induce *ex vivo* filopodia formation, we incubated either control or polyP-enriched GUVs with the minimal number of purified proteins that are necessary to initiate and elongate filopodia *in vitro*, i.e., IRSp53, VASP, β-actin, fascin, profilin, and capping protein^42^. In contrast to the control GUVs where we did not observe a single inward protrusion, polyP-enriched GUVs contained either one or more long filamentous protrusions (Fig. 4K, L). Based on these results, we propose that membrane-associated polyP acts as a scaffolding nexus for IRSp53 which positively affects filopodia numbers and lengths in mammalian cells.

### PolyP levels correlate directly with adhesive capacity and inversely with cell migration

Filopodia continuously assemble and disassemble as they probe their environment ^26,52^. Whereas more stable filopodia promote tight adhesion and slow migration, more dynamic filopodia lead to the opposite effects^28,29,53,54^. Based on our earlier results, we now wondered whether changes in cellular polyP levels affect a cell’s decision between adhesion and migration. Analysis of the migratory and adhesive capacity of mock-, EcPPK1-, or ScPPX-transfected NIH3T3 fibroblasts revealed that cells with higher-than-normal levels of polyP (EcPPK1-NIH3T3) showed an overall decrease in migration across a porous membrane (Fig. 5A,B) and a corresponding increase in adherence (Fig. 5C) compared to mock-transfected control cells. In contrast, cells with depleted polyP levels (i.e., ScPPX-NIH3T3 plus Shield) showed higher trans-well mobility (Fig. 5D, E) and lower adherence capacity (Fig. 5F) than control cells with unaltered polyP levels (*i.e.*, ScPPX-NIH3T3 minus Shield). We independently confirmed some of these results in U2OS (Fig. S5A, B) and HeLa (Fig. S5D, E) cells. These results suggested that alterations in cellular polyP levels affect the migratory behavior of cells; while raising polyP levels increases cell adhesion, lowering cellular polyP levels appears to increase cell migration.

**Figure 5.**
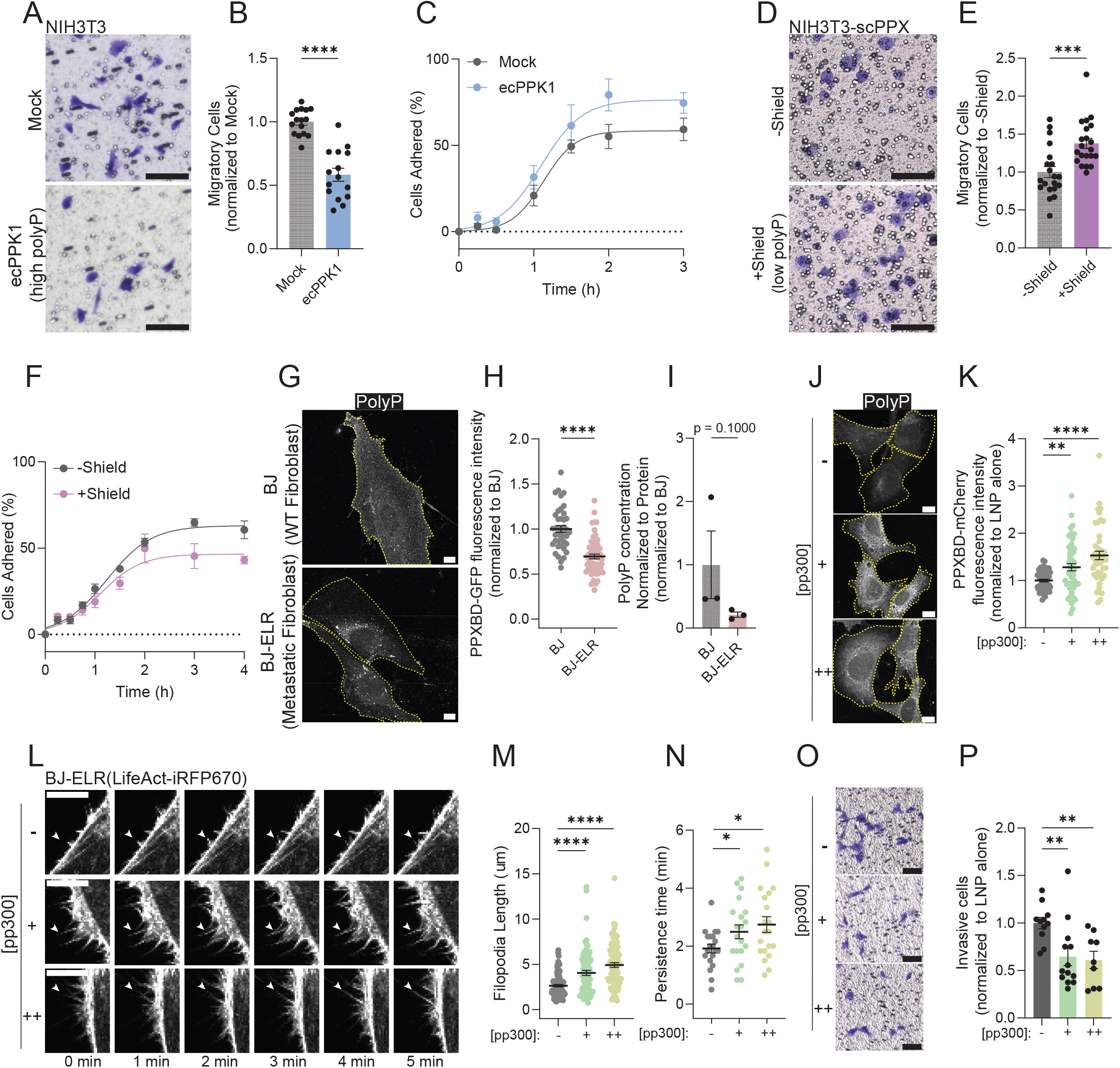
PolyP promotes cell adhesion and reduces trans-well migration. (**A**) Trans-well migration of mock- or EcPPK1-transfected NIH3T3 fibroblasts in a Boyden chamber. After 4 hours, the migrated cells were fixed and stained with crystal violet (CV). (**B**) Quantification of data shown in (**A**). Each data point represents the total number of CV-stained cells in a randomly selected field of view. Three distinct fields of view were analyzed per condition and replicate (n=3). Number of migrating mock-transfected cells was set to 1. (**C**) Time course of mock- or EcPPK1-transfected NIH3T3 fibroblasts adherence on glass bottomed 96 well-plates. Adhered cells were fixed, stained with CV and measure for absorbance. Each data point represents the average percentage of adherent cells in three biological replicates. CV absorbance from a well in which cells were allowed to adhere for 24h was set to a 100%. (**D**) Trans-well migration of NIH3T3-DD-3xFLAG-ScPPX fibroblasts pretreated with or without the stabilizer Shield for 24h as described in (A). (**E**) Quantification of the trans-well migration data shown in (**D**). (**F**) Time course of NIH3T3-DD-3xFLAG-ScPPX fibroblasts pre-treated with or without the stabilizer Shield for 24h as described in (**C**). (**G**) BJ (WT) and BJ-ELR (metastatic) human fibroblasts stained with PPXBD-GFP to visualize cellular polyP levels. See Fig. S5F for biological replicates. (**H**) Quantification of the fluorescent polyP signal shown in (**G**). Each data point represents the mean PPXBD-GFP fluorescence intensity of a single cell (n=15 per condition and replicate; n=3). Average fluorescence intensity of BJ fibroblasts was set to 1 for each replicate. (**I**) Quantification of extracted polyP. Each data point represents polyP levels in ∼200,000 cells; n=3. (**J**) BJ-ELR fibroblasts were incubated with buffer-loaded (-) or polyP-loaded (+:10 mM; ++:50 mM polyP-300) LNPs for 24 h and stained with PPXBD-GFP to visualize cellular polyP levels. See Fig. S5G for biological replicates. (**K**) Quantification of the data shown in (**J**). See panel (**H**) for details. (**L**) BJELR-LifeAct-iRF670 fibroblasts pre-treated with buffer- or polyP-loaded LNPs for 24h were live-imaged over 5 min. Still images of movie S4 are shown. White arrowheads depict individual filopodia. (**M**) Quantification of filopodia length in cells shown in (**L**). Each data point represents the length of a single filopodium; n=3 cells. (**N**) Persistence times of similarly-length filopodia of cells shown in (**L**). Each data point represents a single filopodium tracked over time with a maximum of 3 filopodia arising from a single cell; n=3. (**O**) Invasion of BJELR fibroblasts pre-treated with buffer-or polyP-loaded LNPs for 24h using a collagen-coated Boyden chamber. After 24h, the invaded cells were fixed and stained with CV. (**P**) Quantification of the trans-well invasion data shown in (**O**). See panel D for details. Scale bars: 100 µm for Boyden chamber assays; 10 µm for IF images. Unpaired t-tests were used to compare pairs of samples in B, E, H and I whereas one way ANOVA was used to analyze samples with multiple comparisons in K, M, N and P with a p-value threshold of 0.05. Quantitative image analysis for panels B, E, H, K, M, N, was conducted blinded.

### Metastatic fibroblasts show a significant reduction in polyP levels

Previous studies revealed that polyP levels in patient-derived cancer cells correlate inversely with their sensitivity towards cisplatin-induced apoptosis, with cancer cell lines showing the highest resistance to the anticancer drug containing the lowest levels of polyP^4^. Increasing the endogenous polyP levels resensitized these cancer cells towards cisplatin, suggesting that cancer cells might benefit from downregulating their polyP levels. To test whether cancer cells undergo a reorganization of their cellular polyP pool, which, based on our current study, would dysregulate filopodia formation and potentially lead to increased invasion, a hallmark of metastatic transformation of cells^53,55–57^, we compared endogenous polyP levels in the naïve human BJ cells with an isogenic metastatic BJ-ELR cell line^58^. These cells, which show a significantly higher degree of invasion as assessed by the trans-well invasion assay compared to BJ cells (Fig. S5E, F), were generated by transforming BJ cells with the telomerase catalytic subunit (*hTERT)*, the simian virus 40 large-T oncoprotein and the oncogenic allele of H-*ras*^58^. Analysis of the endogenous polyP signal revealed an almost 50% downregulation of total cellular polyP levels in BJ-ELR cells (Fig. 5G-I), with most of the remaining polyP pool located close to or within the nucleus (Fig. 5G, S5G). This result was also consistent with previous studies in HeLa cells, where most of the polyP signal was found associated with the nucleus^4^. These results suggested that some cancer cells undergo a significant polyP reorganization, depleting and/or re-distributing their membrane-associated polyP. We now reasoned that if the observed reorganization of polyP was indeed associated with increased cell migration and invasion in cancer cells, we should be able to reverse these phenotypes by restoring the cellular polyP levels. However, expression of EcPPK1 in mammalian cells, while useful for transiently increasing polyP levels and monitoring the physiological outcome, is not suitable for long-term experiments because even leaky expression of EcPPK1 eventually becomes cytotoxic^18^. To avoid this issue, we tested the possibility of delivering polyP-300 via commercially available lipid nanoparticles (LNP). We considered this approach to be superior to the simple incubation of mammalian cells with polyP because prior studies revealed that addition of polyP to the media can trigger the further depletion of endogenous polyP stores^59^. To investigate whether LNP-mediated delivery increases the cytosolic levels of polyP, we incubated BJ-ELR cells with two different doses of polyP-300 (pp300)-enriched LNPs for 24h (+, 10 mM polyP-300; ++, 50 mM polyP-300). Image-based quantification confirmed a significant increase in cellular polyP levels (Fig. 5J,K). In full agreement with our previous results, live-cell imaging of Life-Act IRFP670-transfected BJ-ELR demonstrated that cells exposed to LNP-mediated delivery of polyP-300 produced significantly longer (Fig. 5M) and more persistent (Fig. 5N) filopodia than the Life-Act IRFP670-transfected BJ-ELR cells treated with buffer-loaded LNPs. Moreover, and even more crucially, LNP-pp300 treated BJ-ELR cells were significantly less invasive than their control-treated counterparts (Fig. 5O,P). These results suggested not only that cells undergo a dramatic reorganization of the cellular polyP pool upon their transformation into invasive cancer cells but that this change in polyP levels and localization might directly contribute to their increased migratory behavior.

### Restoration of polyP levels reduces invasiveness of breast cancer organoids

To test the hypothesis that restoring levels of polyP in metastatic cancers might reduce their invasive properties, we investigated the effects of LNP-mediated polyP supplementation on the 3D-invasiveness of primary tumor organoids isolated from two different genetically engineered mouse models. In the MMTV-PyMT model, the mouse mammary tumor virus (MMTV) long terminal repeat drives the expression of the polyoma virus middle T oncogene (PyMT), causing the development of a highly aggressive mammary tumor, whose gene expression profile clusters with the aggressive luminal B subtype of human breast cancers^60^. As a second model, we utilized the C3(1)-Tag mouse, a tumor model that represents the highly metastatic basal-like triple-negative breast cancer^61,62^. Organoids prepared from either of these tumors and embedded into fibroblast growth factor (FGF)-supplemented collagen I display robust 3D invasiveness, defined by cells migrating away from the core of the organoid over a 5 to 7-day period, as compared to organoids from healthy mammary tissue ^61,63^. To first determine whether polyP levels and/or distribution differ between healthy mammary tissue and the tumor tissues, we embedded the organoids in collagen I for 24h, prepared cryosections, and stained for polyP. Compared to organoids prepared from healthy murine mammary tissue, the organoids derived from the MMTV-PyMT tumor model (Fig. 6A,B; S6A) or the C3(1)-Tag mouse model (Fig. S6B, C) revealed a significant downregulation of endogenous polyP levels. The most pronounced differences in polyP levels were found along the plasma membrane, where cells in the control organoids show a discrete polyP enrichment that is largely absent in the tumor cells (Fig. 6A; S6A, B). These results provide independent confirmation of our previous results that cancer cells reduce and reorganize their cellular polyP pools. As observed with our BJ-ELR cells, we also found that incubation of the MMTV-PyMT-derived tumor organoids with polyP-300 loaded LNPs for 24h was sufficient to significantly increase their endogenous polyP concentration compared to the organoids from the same tumor that only received buffer-loaded LNPs (Fig. 6C, D; S6D). We then embedded the LNP-treated organoids in collagen I, let them establish for 24h (D0) and added FGF, a key signaling component that induces invasion of tumor organoids into the surrounding collagen gel l^63^. After 7 days (D7), we measured the inverse circularity of the organoids, a readout of their invasive capacity ^64^. We observed that the treatment of the MMTV-PyMT-derived tumor organoids with either of the two polyP300-loaded LNPs significantly (>50%) reduced their invasiveness (Fig. 6E, F; S6E). While the buffer-LNP-treated organoids showed the previously observed irregular growth that extends into the surrounding collagen, the majority of LNP-pp300-treated MMTV-PyMT-derived tumor organoids remained circular. We obtained very similar results when we treated the tumor organoids from C3(1)-Tag-mice with control or polyP-loaded LNPs (Fig. S6F, G), indicating that these effects are not specific to luminal breast cancers. These results suggested that replenishing the cellular polyP pools might be a potential strategy to reduce the invasiveness of metastatic breast cancers.

**Figure 6.**
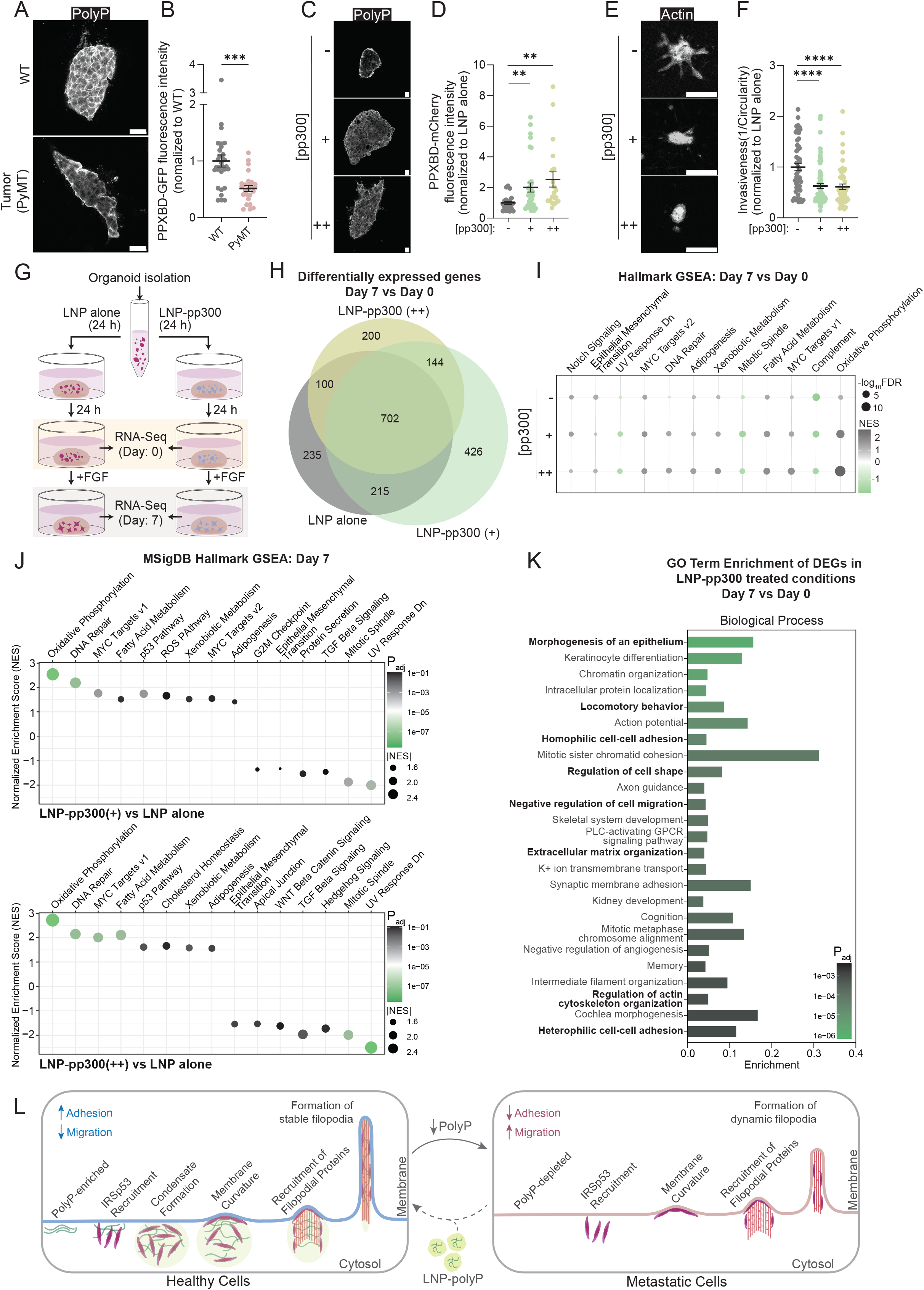
Restoration of polyP levels in organoids reverses metastatic features. (**A**) Fluorescent images of mammary tissue-derived or MMTV-PyMT tumor-derived organoids one day after seeding in collagen gels. Staining with PPXBD-GFP was used to visualize polyP (gray). n=3 mice. Additional images in Fig. S6A. (**B**) Quantification of images shown in (**A**). (**C**) Fluorescence images of MMTV-PyMT tumor-derived organoids pre-treated for 24h with buffer-loaded (-) or polyP-loaded (+,10 mM polyP-300; ++, 50 mM polyP-300) LNPs before seeding in collagen gels. Cryosections were prepared after 24h (D0) and stained for polyP. n=3 mice. Additional images in Fig. S6B. (**D**) Quantification of images shown in (**C**). Each data point in (B, D) represents the average PPXBD-GFP fluorescence intensity of a single organoid. Average fluorescence intensity of the healthy WT (B) or control-LNP (D) organoids was set to 1 per biological replicate. Scale bars (A,C): 10 µm. (**E**) 3D-invasiveness of MMTV-PyMT tumor-derived organoids pre-treated for 24h with buffer-loaded (-) or polyP-loaded (+, 10 mM polyP-300; ++, 50 mM polyP-300) LNPs and allowed to invade for 7 days. The organoids were fixed and stained for DAPI and actin. n=3 mice. Additional images in Fig. S6C. (**F**) Inverse circularity (IC) of organoids shown in (**E**). Each data point represents IC of a single organoid. Average IC of organoids treated with buffer loaded (-) LNPs organoids is set to 1; n=3 mice. (**G**) Experimental outline for RNA extraction and sequencing; n= 4 mice per treatment and time point. Scale bar: 100 µm. (**H**) Venn diagram of differentially expressed genes (DEGs) between D0 and D7 MMTV-PyMT-derived organoids pretreated with buffer-loaded (-) or polyP-loaded (+, 10 mM polyP-300; ++, 50 mM polyP-300) LNPs. (**I**) Dot plot representing the MSigDB Hallmark GSEA pathway enrichment in MMTV-PyMT-derived D7 *vs* D0 organoids under each treatment condition. **(J)** Dot plot representing the MSigDB Hallmark pathway enrichment in D7 MMTV-PyMT-derived organoids treated with either polyP-loaded (+) or polyP-loaded (++) LNPs *vs* buffer-loaded (-) LNPs. (**K**) Gene Ontology (GO) enrichment analysis of DEGs present in D7 *vs* D0 organoids pretreated with LNP-pp300 (+, ++) but not in LNP-control organoids. **(L)** Current working model: Membrane-enriched polyP recruits IRSp53 to form and stabilize mature filopodia which leads to increased adhesion and reduced migration. Cells lose their polyP upon metastatic transformation causing them to form shorter and more dynamic filopodia that are pro-migratory. Replenishing the cellular polyP levels partially alleviates this phenotype.

### PolyP treatment suppresses select invasive gene expression programs

Cancer cells change their transcriptomic profiles as they transition from pre-invasive (D0) to invasive states (D7) ^65^. To test whether polyP treatment affects these transcriptomic changes, we performed RNA-sequencing on D0 and D7 organoids prepared from polyP-treated and control MMTV-PyMT mice (n=4 per treatment and time point) (Fig. 6G). Principal component analysis (PCA) of the normalized D0 and D7 gene expression datasets revealed that the temporal progression was the primary driver of variance, separating D0 profiles from D7 profiles across all subjects and treatments (Fig. S6H). D0 samples indicated significant baseline biological variation between animals. By D7, the samples converged into two distinct sub-clusters. Notably, across all conditions and time points, the LNP alone, LNP-pp300 (+), and LNP-pp300 (++) groups clustered together within each individual animal, demonstrating high technical reproducibility but also suggesting that the polyP-300 treatment did not induce major global shifts in the overall transcriptomic profile compared to the LNP control. In support of this data, we found between 1,200 and 1,400 differentially expressed genes (DEGs) across the 7-day time-period (D7 *vs* D0) for each condition, with the majority of gene expression changes shared between the polyP treatments and control (Fig. 6H; Table S1). Nonetheless, gene set enrichment analysis (GSEA) revealed a number of pathways that were commonly up and downregulated for all three conditions, however, several cancer hallmark pathways displayed different levels of enrichment upon polyP treatment (Fig. 6I). The largest increase was observed in oxidative phosphorylation enrichment upon polyP treatment (Fig. 6I; S6I). This result was fully consistent with previous studies that revealed that OXPHOS is stimulated by polyP ^66^. Other significantly more enriched hallmark pathways in polyP-supplemented D7 *vs* D0 organoids included fatty acid and xenobiotic metabolism, both likely connected to the observed increase in OXPHOS. On the flip side, one of the more significantly depleted pathways in polyP-supplemented organoids compared to control organoids was the epithelial to mesenchymal transition (EMT) pathway, a major driver of metastasis ^67^. Analysis of the genes underlying the suppression of this pathway, included changes in several actin-related genes (*e.g.,* Myl9, Acta2) as well as genes encoding for ECM and adhesion proteins (*e.g.,* Postn, Sntb1) and transcriptional regulators, known to promote invasion and metastasis (*e.g.,* Snai2) (Fig. S6I).

We also observed the downregulation of the EMT pathway when we directly compared the expression changes between invasive organoids (D7) upon LNP-pp300 and control treatment (Fig. 6J). In addition, we also noted a significant downregulation of several other pro-metastatic pathways including apical junction, WNT signaling, G2M checkpoint and mitotic spindle pathways upon polyP-treatment (Fig. 6J). Moreover, this comparison also confirmed the significant upregulation of the OXPHOS, p53 and ROS pathways, all three of which typically associated with lower tumor aggressiveness ^67–69^. Finally, analysis of the biological processes associated with polyP-specific gene expression changes upon invasion revealed a significant enrichment for genes involved in the negative regulation of cell migration and cell-cell adhesion (Fig. 6K). These results provide independent support for our conclusion that polyP serves as a regulator of pro-adhesion and anti-migration.

## Discussion

### PolyP – A primordial protein scaffold

PolyP emerged with the formation of the Earth’s crust several billion years ago^70^. Equivalent to ATP in chemical energy yet entirely inorganic in nature, it has been hypothesized to have played a pivotal role in the formation of the germinal progenote, the earliest speculated cell-like system preceding cellular life^71^. Conserved throughout evolution, polyP remains present in all major subcellular compartments across highly diverse organisms. In mammals, its synthesis is likely ATP-based yet the precise mechanism remains still largely enigmatic^10^. PolyP’s functional versatility appears unprecedented and, dependent on its subcellular localization, it is involved in rRNA transcription, calcium homeostasis, energy metabolism, cell signaling, stress responses and amyloid formation ^2,5^. The central unifying feature of polyP’s seemingly unrelated functions appears to be its ability to interact with proteins in various conformational states, ranging from stabilizing protein folding intermediates in the form of small, soluble oligomers^72^ and scaffolding amyloidogenic proteins into distinct fibril morphologies^73,74^ to driving RNA/DNA and, based on our studies here, potentially membrane-binding proteins into phase-separated liquid droplets^75,76^. This idea is fully consistent with a model in which primordial polypeptides evolved in a polyP-rich environment and gained selective advantage by utilizing polyP’s scaffolding capacity for their individual functional and structural optimization. Here we followed up on earlier studies, which revealed that some of the highest concentrations of polyP in primary cells are found along the plasma membrane. We now demonstrate that membrane-associated polyP aids in the formation and stabilization of filopodia and reveal that cancer cells, by depleting and reorganizing their cytosolic polyP pool, exploit this function to make cell adhesion *versus* migration and invasion decisions. We furthermore present evidence that targeted re-supplementation of polyP might have anticancer treatment potential.

### PolyP – A regulator of filopodia dynamics

Filopodia are well-characterized actin-rich membrane protrusions, which play critical roles in cell-cell adhesion in epithelial sheets, dendrite formation in neurons, and ECM sensing in fibroblasts ^27,77^. The formation of filopodia at the membrane has been described by two distinct models; the convergent elongation model, which proposes that filopodia expand on a pre-formed actin network in the lamellipodium^78^, and the tip nucleation model, which postulates that filopodia formation starts *de novo* by recruiting filopodial initiation proteins, particularly IRSp53, to the inner leaflet of membranes, locally enriched in PIP2^79,80^. Common to both models is the requirement for a negative curvature in the plasma membrane, which is promoted by the multimerization of activated IRSp53^32,40,42^. A recent *in silico* study proposed that IRSp53 dimers are on their own rather straight and only induce a high degree of negative curvature upon stacking into higher-order oligomeric structures^47^. Our results now suggest that membrane associated polyP might play a role in this process by assisting in the scaffolding of membrane-associated IRSp53 dimers. We base this conclusion on several independent findings; i) membrane-associated polyP colocalizes with both IRSp53 and CDC42, the two critical components that initiate filopodia formation; ii) membrane-associated polyP is sufficient to recruit purified IRSp53 into distinct clusters along GUV membranes, promoting filopodia formation *ex vivo*; and iii) polyP induces the phase separation of IRSp53, an assembly mechanism that has been proposed to contribute to IRSp53’s functional properties on the membrane^48,81^. The latter is especially relevant for cortical neurons, where the phase separation of IRSp53 along with its binding partners PSD-95 and Shank3, appears to be required for appropriate synapse formation and function^48^. We postulate that the interaction between IRSp53 and long-chain polyP promotes its clustering and multimerization through phase separation, a critical step in inducing membrane curvature and protein recruitment (Fig. 6L). Of note, IRSp53 is not only involved in the initiation of filopodia but also tethers actin filaments to the plasma membrane ^82^. Based on the colocalization of polyP and IRSp53 along the filopodia axis as well as our cryo-ET data, which show a significantly denser protein network connecting the filopodial membrane and actin filaments in EcPPK1-expressing cells compared to non-transfected cells, we argue that polyP uses a similar mechanism along the filopodial axis to promote IRSp53-actin interactions, reduce filopodia dynamics and increase their length, number and likely adhesive properties.

### The PIP2-polyP connection

One intriguing question that has long concerned the polyP-field regards the mechanism by which polyP, itself a highly negatively charged molecule, is able to accumulate at distinct regions of the plasma membrane ^9^. Our discovery that polyP interacts with membrane-associated IRSp53 at filopodial initiation sites might finally yield an answer. Previous studies demonstrated that IRSp53 is recruited to membrane regions that are specifically enriched in PIP2^42^, a precursor of inositol phosphates (IPs). Of note, several different IPs have long been linked to mammalian polyP production^83^. For instance, yeast mutants defective in inositol pyrophosphate synthesis show no detectable polyP^84^ while in mice, deletion of inositol hexakisphosphate kinase 1 (IP6K1), which generates high-energy inositol pyrophosphates like InsP7, leads to a significant reduction in platelet polyP levels and a loss of polyP-mediated functions in blood clotting^85^. The direct mechanism by which these PIs and polyP influence each other, however, remains unknown. Because it is not known how polyP chains are formed *de novo,* it has been suggested that IPs might serve as the central “seed” for growing polyP chains ^83,85^. This mechanism would not only explain the presence of polyP on PIP2 enriched membranes but would suggest that some of the previously conducted mechanistic studies on the role of PIP2 in filopodia formation ^42,86^ might need to be revisited in the context of polyP.

### Reorganization of endogenous polyP – A new molecular hallmark of cancer cells?

Cellular mechanisms that control cell adhesion and filopodia formation are known to become dysregulated during the metastatic transformation of cells^57^, trading filopodia-mediated cell-cell and cell-surface adhesion with a significant increase in lamellipodia^87^. Yet the molecular mechanisms of tumor migration and invasion are dependent on the type of cancer and hence the cell type of origin^88^. In our study, we use mouse models of two types of highly aggressive breast cancers, a luminal B type and a basal-like triple-negative breast cancer (TNBC)^60,61^. These subtypes of cancers are prominently characterized by their aggressive remodeling of the actin cytoskeleton, which drives decreased filopodial stability and increased invasive capacity^89,90^. Key filopodial proteins that are differentially regulated in these cancers include WAVE-3, whose upregulation increases the formation of lamellipodia and lamellipodia-derived filopodia^91^ and formin DIAPH3, whose downregulation increases the expression of certain invadopodial proteins in TNBC^92^. Our work now adds dysregulation of polyP as a new breast cancer signature. We found that cancer cells undergo both a significant overall decrease in polyP levels and a dramatic reorganization of its subcellular distribution (Fig. 6L). Analysis of the current literature revealed that at least two polyphosphatases become significantly upregulated upon metastatic cell transformation; NUDT3, a recently characterized mammalian endopolyphosphatase, which degrades polyP in a zinc dependent manner^93^, and Hprune-1, a mammalian exo-polyphosphatase^12,94,95^. NUDT3 levels are heavily increased in lung adenocarcinomas and TNBCs, the latter ones also known for their uncharacteristically high levels of zinc ^96,97,98^. Dependency map (Depmap) analysis confirmed this strong upregulation of both NUDT3 and H-Prune1 expression in highly aggressive forms of breast cancer^99^, suggesting that the observed downregulation of polyP might be part of a program that transforms cells into metastatic cancer cells. Of note, the oncogenic potential of NUDT3 has been historically ascribed to its mRNA decapping activity, which has been proposed to alter cell migration ^98^ while upregulation of H-Prune1 has also been shown to promote cell motility and increased invasion ^99^. Our data now demonstrate that the loss of membrane-bound polyP might be the critical factor that causes many of these observed effects. Moreover, the decrease of cellular polyP levels in cancer cells likely contribute to other critical processes that mediate cell proliferation; among the many reported functions of polyP that might be relevant in this context are its reported inhibitory effect on RNA-Pol2 activity^17,18^, its stimulatory effect on mitochondrial bioenergetics^20^, and its pro-apoptotic activity in cancer cells^4^. These would lead to an increase protein translation and likely growth rates, contribute to the reported metabolic switch from OXPHOS to glycolysis, and increase resistance to select anti-cancer treatments, respectively ^100,101^. Future studies in other types of cancer are needed to test the idea that the reorganization of one of the most conserved and multifactorial molecules in eukaryotic cells might be a new molecular hallmark of cancer.

### Replenishing polyP stores in cancer cells reverses metastatic features of cancer cells

Earlier studies from our lab conducted in HeLa and patient-derived ovarian cancer cells demonstrated that low endogenous polyP accompanies resistance to apoptosis and to chemotherapeutics such as cisplatin^4^. Moreover, raising polyP levels in solid colon tumors has been shown to inhibit their growth in mice^102^. Two recent colon cancer studies, however, reported the opposite effect: by exogenously administering polyP, the authors observed increased tumor proliferation^103,104^. Whether this discrepancy is cancer-type-specific or a consequence of the delivery route remains to be tested, particularly as exogenously administered polyP can paradoxically trigger polyP release, depleting the very endogenous pools it is meant to raise^59^. Our delivery strategy avoids this confounding issue; we demonstrated that LNP-mediated polyP delivery restores the depleted polyP stores, a feature that characterizes a number of different cancer cells. We found that this treatment reduces cell migration and suppresses the 3D invasiveness of metastatic tumor organoids. Transcriptomic analysis revealed three coherent classes of changes: pathways previously linked to polyP function, such as OXPHOS, changed in the expected direction; transcriptional programs that track with aggressive disease, including EMT and apical-junction disorganization, shifted toward a less malignant state; and processes governing cell migration and adhesion moved toward reduced motility and stronger adhesion. The migration and adhesion effects most likely follow directly from restored membrane-associated polyP, whereas the broader pathway remodeling probably reflects polyP’s compartment-specific functions elsewhere in the cell. Because maintaining endogenous polyP pools appears to prevent many of the cellular changes that accompany metastasis, we propose that polyP serves as an evolutionarily conserved brake on cell proliferation, making its downregulation a driver of metastasis (Fig. 6L). Restoring endogenous polyP might thus be a powerful new therapeutic strategy to treat cancer metastasis.

## LIMITATIONS OF THE STUDY

Some of the questions that remain to be solved include how polyP drives IRSp53 oligomerization and phase separation. A detailed structural investigation will be needed to identify and subsequently manipulate the critical polyP–IRSp53 interaction sites while leaving IRSp53 itself intact; expressing a variant engineered at that interface would ultimately reveal whether polyP-mediated phase separation of IRSp53 is genuinely required for filopodia formation. We also have not established whether or by what mechanism polyP is anchored at the membrane, nor the precise role of inositol phosphates (IPs) in that process. This is not an idle gap, given that enzymes of IP synthesis affect many downstream processes, which makes data interpretation very challenging. Finally, before proposing that polyP depletion is a *unifying* feature of metastatic cancers, a systematic survey across a broad set of patient-derived organoids is required, measuring polyP against non-tumor tissues and tracking invasive behavior after LNP-mediated polyP supplementation.

## RESOURCE AVAILABILITY

### Lead contact

Further information and requests for resources and reagents should be directed to and will be fulfilled by the lead contact, Ursula Jakob.

### Materials availability

All cell lines generated in this study are available upon request from the lead contact with a completed materials transfer agreement.

### Data and code availability

Unprocessed blots, gels, and microscopy images are available at Mendeley Data DOI:10.17632/3bvc4vj8ft.1. The GEO accession code is GSE344396. Any additional information required to reanalyze the data reported in this paper is available from the lead contact upon request.

## ACKNOWLEDGEMENTS

We thank Michael Downey and Brent Stockwell for providing us with some of the vectors and cell lines. We thank T. Shiba (RegenTiss, Japan) for purified polyP. We thank Aileen Ariosa and Jeff Johnson from Cayman Chemicals for providing us with LNPs. We thank Ken Wan for purifying the proteins and Jim Bardwell, Morgan DeSantis, and Ann Miller for many helpful discussions. This work was supported by the NIH grants GM122506 to U.J., DP2GM150019 to S.M., GM163198 to A.L. and the Emerald Foundation Early Career Award to J.W. Research reported in this publication was supported by the NIH S10OD030275 and the Arnold and Mabel Beckmann Foundation grants to the University of Michigan Cryo-EM Facility (U-M Cryo-EM). U-M Cryo-EM is grateful for support from the U-M Life Sciences Institute and the U-M Biosciences Initiative.

## AUTHOR CONTRIBUTIONS

**A.R.** Conceptualization, Methodology, Writing- Original draft preparation; Validation; Formal analysis, Investigation, Visualization, Supervision; **A.J.** Investigation, Validation, Formal analysis; **H.Z.** Investigation, Formal analysis; **J.G.** Investigation, Visualization, Formal analysis; **P.M.** Investigation, Formal analysis; **B.J.O.** Visualization; **M.A.** Investigation, Resources; **Y.X.** Investigation, Resources; **J.S.** Investigation, Formal analysis; **M.Su.** Investigation, Formal analysis; **H.R.** Resources; **B.O.** Visualization, Formal analysis; **A.E.** Investigation, Visualization, Formal analysis; **A.B.J. B.** Investigation, Formal analysis; **J.O.S.** Validation, Formal analysis; **A.L.** Supervision, Project administration, Funding acquisition; **M.S.** Supervision, Funding acquisition, Editing; **J.W.** Supervision, Project administration, Funding acquisition, Editing; **U.J.** Conceptualization, Supervision, Reviewing and Editing, Project administration, Funding acquisition.

## MATERIAL AND METHODS

### Cell Lines and Culture

NIH3T3 (ATCC CRL-1658™), COS7 (ATCC CRL-1651™), HeLa (ATCC CCL-2™), U2OS (ATCC HTB-96™), BJ fibroblasts (ATCC CRL-2522™) and BJ-ELR transformed fibroblasts (gift from Dr. Brent Stockwell, Columbia University, New York) were grown and maintained in DMEM (Gibco™ 11995065) supplemented with 10% w/v Fetal Bovine Serum (FBS) (#F4135, Sigma-Aldrich) and 1% w/v Penicillin-Streptomycin (#SV30010, Cytiva). All cell lines were maintained at 37°C and 5% CO_2_. The cell lines were regularly tested against mycoplasma.

### Plasmids, transfections and stable cell line generation

The EcPPK1 plasmid (Addgene plasmid # 108850) was a gift from Michael Downey. Transient transfections were carried out on cells seeded on glass coverslips in a 24-well plate using Lipofectamine LTX Reagent with PLUS Reagent (15338100, ThermoFisher Scientific) according to the protocol provided on the manufacturer’s website. The N-terminally 3xFLAG-tagged ScPPX gene was cloned into the pLVX-pTuner Green vector (#632176, Takara Bio). This vector contains the coding sequence for an N-terminal destabilization domain (DD), which leads to the degradation of the protein in the absence of the small molecule Shield1 (#632189, Takara Bio). For stable transfections, lentiviruses were generated encoding this construct (University of Michigan Vector Core). NIH3T3 cells were transfected with the DD-3x-FLAG-ScPPX lentivirus in the presence of polybrene (#TR-1003-G, EMD Millipore) at a concentration of 10 µg/ml for 24 hours. Transfection media was removed and cells were allowed to recover for 24 hours. Transfected clones were sorted based on positive ZsGreen fluorescence (B525 channel) using the Cytoflex SRT single cell sorter (Beckman Coulter). The sorted cells were grown in the presence or absence of 0.5 µM Shield1. The expression of 3xFLAG-ScPPX was verified by western Blotting using a mouse anti-FLAG M2 antibody (#F3165, Sigma-Aldrich). To generate stably transfected LifeAct-iRFP670 expressing cell lines, lentiviruses were generated as before using the pLViP-LifeAct-iRFP670 construct (Addgene plasmid # 229699). NIH3T3, NIH3T3-ScPPX and BJ-ELR cells were transfected with the lentivirus in the presence of polybrene (#TR-1003-G, EMD Millipore) at a concentration of 10 µg/ml. Cells were transfected for 24 hours and allowed to recover for 24 hours after removal of the transfection media. Positively transfected clones were sorted based on iRFP670 fluorescence (R660 channel) using the Cytoflex SRT (Beckman Coulter). Monoclonal populations were prepared from all lentivirus-transfected cell lines by growing individual sorted cells in conditioned media from the respective cell lines, supplemented with 20% w/v FBS in 96-well plates.

### Immunofluorescence staining of fibroblasts and image analysis

To image mammalian cells by fluorescence microscopy, cells were detached using 0.25 % Trypsin-EDTA (T4049, Sigma-Aldrich) and seeded onto 12 mm circular coverslips (#72230-01, Electron Microscopy Sciences) at a density of 25,000 cells/well and placed into a 24-well plate. Following transfections and/or the treatment of cells seeded on coverslips, cells were fixed with freshly prepared 4% v/v paraformaldehyde (1578100, Electron Microscopy Sciences) for 20 mins. Fixed cells were washed three times with 1X phosphate-buffered saline (PBS) and permeabilized with PBS solution containing 0.3% v/v Triton X-100 (T8787, Sigma-Aldrich) and 1% w/v bovine serum albumin (BSA) (A3059, Sigma-Aldrich) for 1 hour. After the permeabilization, the cells were washed with 1xPBS and incubated in a solution of 1% w/v BSA in PBS (blocking solution) for 1 hour. To visualize endogenous polyP, cells were incubated with 10 μg/ml purified PPXBD-eGFP^4^ or, in the case of ScPPX-transfected cells, PPXBD-mCherry^4^, in blocking solution overnight at room temperature. Cells were co-stained with primary antibodies (all antibodies listed in Key Resources Table) using the recommended dilution overnight at 4°C. Cells were washed 3 times in 1x PBS prior to the addition of a 1:1000 dilution of the secondary antibodies in blocking buffer. After 2 hours of incubation at RT, the cells were washed with 1x PBS and incubated with a mix of 1.32 µM actin-stain Phalloidin-647 (#8940, Cell Signaling Technology) and 1 μg/ml DAPI (#D1306, Thermo Fisher Scientific) for 15 mins. The coverslips were washed three times before mounting them on a microscope objective slide using Prolong Gold (9071S, Cell Signaling Technology). After sealing them with nail polish, images were taken after at least 24 hours. The cells were visualized using a 63x/100x oil objective on a Leica SP8 laser scanning confocal (Leica GmbH, Mannheim Germany) on a DMI8 base microscope (or) on a Leica Stellaris microscope with a 63x oil immersion objective (Leica GmbH, Mannheim Germany) using the LAS X software. Analysis of fluorescence intensities was done by using the actin stain to obtain ROIs of individual cells and the mean signal intensity values of the resulting ROIs were collected and plotted on Graphpad Prism 11 version 11.0.2. The average filopodia count was obtained by counting the total number of filopodia (i.e., linear F-actin rich protrusion tipped with MyoX) in a field of view and dividing them by the total number of cells as quantified by DAPI staining in the same field of view. A minimum of 30 cells was analyzed per condition and replicate using FIJI. All images were blinded prior to analysis. Filopodia lengths were calculated by drawing a segmented line from the base of the filopodia to the tip using the FIJI software. A minimum of 30 filopodia from 5-10 different cells were analyzed per condition and replicate.

### Live cell fluorescent imaging and image analysis

Imaging of the fixed cells was performed using a 63x/100x oil objective on a Leica SP8 laser scanning confocal (Leica GmbH, Mannheim Germany) on a DMI8 base microscope (or) on a Leica Stellaris microscope with a 63x oil immersion objective (Leica GmbH, Mannheim Germany) using the LAS X software. Live cell imaging was performed on a Leica SP8 inverted microscope equipped with an incubated chamber (Tokai) maintained at 37°C and 5% CO_2_ using a 63x oil immersion objective and supported by the LAS X software (Leica GmbH, Mannheim, Germany). Images were acquired every 10 seconds for 5 mins to observe filopodia formation. To measure latrunculin B sensitivity, ∼10,000 cells were seeded in an 8 well chamber slide and transfected with mock- or EcPPK1. 24 hours post transfection cells were live-imaged as before except that images were acquired every 15 seconds. After 10 min, 250 nM latrunculin B was added and cells were imaged every 15 seconds for another 20 mins to measure filopodia retraction. The persistence times of filopodia in the absence and presence of latrunculin were calculated using the manual tracking plugin of the FIJI software, which tracks the number of frames that it takes for fully formed filopodia of equal length to completely retract. The persistence times were graphed using GraphPad Prism version 11.0.2.

### Polyphosphate extraction and quantification

The protocol to extract and quantify polyP was adapted from^18^. Briefly, ∼200,000 cells were trypsinized and washed twice in PBS before finally pelleting the cells and removing all the supernatant. The cell pellet was flash frozen and stored in liquid N_2_ until further use. To lyse the cells, the pellet was thawed on ice. Cells were resuspended in 500 µl 1 M perchloric acid and incubated at RT for 10 mins under constant shaking followed by a 1-min centrifugation at 2,400xg. The supernatant was transferred to a tube containing 4 mg titanium oxide beads (Titansphere 5 µm, 500 mg, #5020-75000, GL Sciences). PolyP was allowed to bind for 20 mins at 4°C with constant rotation. After pelleting the beads, the lysate was discarded and the beads were washed twice with 500 µl 0.1 M perchloric acid before their final resuspension in 200 µl ice-cold 2.8 % v/v ammonium hydroxide solution to release the bound polyP. The beads were vortexed briefly and incubated for 5 mins at 4°C with constant shaking. The beads were pelleted by centrifugation at 2,400xg for 1 min and the supernatant was transferred into a new tube, evaporated completely using a Rotavap and resuspended in 20-40 µl ddH_2_O. The amount of polyP in the sample was determined using a fluorometric DAPI binding assay^105^ at 415/550 nm (ex/em) and a polyP-300 standard curve as reference. The DAPI fluorescence shift was normalized to the total protein concentration in the lysates as determined by SDS-PAGE and densitometry measurements of the Coomassie-stained gels. Samples for each repeat were processed and analyzed at the same time.

### Western blot analysis

Cells were lysed in 1x Laemmli loading buffer and boiled for 5 minutes. Electrophoresis and transfer were conducted as per standard protocols. The membrane was blocked with 5% w/v BSA in 1X-TBST (Tris-Buffered Saline, 0.1% v/v Tween 20). Primary and secondary antibodies were diluted according to the manufacturer in 5% w/v BSA, 1X-TBST and blots were incubated at 4°C overnight or at RT for 45 min, respectively. The bands were visualized using enhanced chemiluminescence (SuperSignal West Pico PLUS, Thermo Fisher Scientific, 34580). The following primary antibodies were used in this study: anti-MyoX, anti-CDC42, anti-BAIAP2/IRSp53, anti-FLAG M2, Anti-fascin. The secondary antibodies used in this study were goat anti-rabbit IgG (H+L)-HRP or goat anti-mouse IgG (H+L)-HRP (Invitrogen). See key resources table for details.

### ATP measurements

The CellTiter-Glo 2.0 Assay kit (#G9241, Promega) was used to determine ATP concentration in the supernatant of cell lysates. Briefly, ∼ 10,000 cells were resuspended in media and mixed at a 1:1 ratio with pre-equilibrated CellTiter-Glo 2.0 reagent and pipetted into an opaque-walled 96-well plate. The plate was incubated at room temperature for 10 mins to allow cell lysis and the signal to stabilize. Luminescence was measured using an Infinite M1000 (Tecan) plate reader. A standard curve using defined ATP concentrations diluted in cell culture media was used to determine the ATP concentration in the cell lysates (#R0441, Thermo Scientific). The same number of cells were lysed and run on a stain-free gel (BioRad). The signal was analyzed using densitometry plots and used to normalize the ATP levels. Samples for each repeat were processed and analyzed at the same time.

### *In situ* cryo-electron tomography of NIH3T3 fibroblasts

#### Grid preparation

Gold grids, 200 mesh, with holey R1/4 SiO_2_ film (Quantifoil Micro Tools GmbH, Q250AR-14S) were glow-discharged for 30 s at 5 mA using an EasiGlow system (Pelco). To achieve cell growth in the center of grid squares, grids were coated with poly-L-lysine (Sigma-Aldrich, P6282) and mPEG-SVA (Laysan Bio, MPEG-SVA-5000). Coated grids were then photosensitized by adding PLPP gel (Alvéole, B004) and photo-micropatterned using a DMi8 microscope (Leica Microsystems) equipped with an Alvéole PRIMO 2 to generate circular patterns with a diameter of 40 µm at the centers of grid squares. Regions of interest were identified, and patterns were selected using the Leonardo 5 photopatterning software (Alvéole). Grids were next coated with 50 µg/mL fibronectin (Sigma-Aldrich, 341631) in a 35 mm glass-bottom dish (MatTek Life Sciences, P35G-1.5-14-C). *Sample preparation:* 1×10^5^ mock or EcPPK1-transfected NIH3T3 fibroblasts were grown in 35 mm glass-bottom dishes containing micropatterned Au SiO_2_ R1/4 200 mesh grids (Quantifoil GmbH) coated with fibronectin (Sigma-Aldrich). Grids were blotted back-side only and plunge-frozen into a liquid propane/ethane mixture (63:37%) at −190°C using a GP2 (Leica Microsystems) automatic grid plunger with the following specifications: blot time: 11 s, temperature: 37°C, and humidity: 80%. The plunge-frozen grids were clipped with C-Clips (Thermo Fisher Scientific, 1036171) into AutoGrids (Thermo Fisher Scientific, 1205101) and stored in liquid nitrogen until further use.

#### Tilt series data acquisition and tomogram reconstruction

Grids containing mouse embryonic fibroblasts were imaged on a 300kV Titan Krios G4i transmission electron microscope (Thermo Fisher Scientific) equipped with a K3 direct electron detector and an imaging filter (Gatan Inc.) operated in counting mode. Dose-symmetric tilt series ^106^ were collected at sites with filipodia from −51° to +51° at 3° intervals, using SerialEM software^107^. The magnification was set to 2.651 Å/pixel, and the total dose per tilt series was 160 e^−^/Å². Data were acquired over a defocus range of −3 µm to −5 µm and with a 20 eV energy-filter slit width. Programs used for tilt series alignment and reconstruction were accessed via SBGrid^108^. A custom script implementing WarpTools (https://github.com/warpem/warp) was used to align and reconstruct tilt series via patch tracking and weighted backprojection, as implemented in IMOD ^109^. Nonlinear anisotropic diffusion (NAD) filtering was applied to reconstructed tomograms in *etomo* ^110^(10 iterations; K=5) for visualization and analysis.

#### Quantification

The number of membrane-associated densities per micron of plasma membrane was determined by manually counting them within the filipodia and dividing by the total membrane length (# membrane-associated densities/µm). The length of the plasma membrane was measured using the measure feature in 3dmod from the IMOD package ^109^. Values were plotted using GraphPad Prism version 11.0.2. A Welch’s t-test was performed using GraphPad Prism version 11.0.2 to assess statistical significance.

### Proximity Ligation Assay (PLA)

PLA was conducted using the Duolink® In Situ Proximity Ligation Assay Starter Kit, Red, Mouse/Rabbit (#DUO92101, Sigma-Aldrich). In brief, NIH3T3 cells were fixed on glass coverslips (#72230-01, Electron Microscopy Sciences) in 4% v/v PFA for 20 min at RT. The fixed cells were washed and permeabilized for 1hr at RT in PBS supplemented with 0.1% v/v Triton X-100, 1% w/v BSS. After incubating the coverslips in 1% w/v BSA (in PBS) for 1 hr at RT, cells were incubated with either 10 µg/ml purified GFP or PPXBD-GFP overnight at 4°C. The next day, the coverslips were rinsed and incubated with mouse anti-GFP (1:100) (sc-9996, Santa Cruz Biotechnology) or mouse anti-beta-actin (LI-COR Biosciences, #926-42212) and rabbit anti-IRSp53 (#HPA023310, Atlas Antibodies) overnight at 4°C. The coverslips were then washed in 1x PBS prior to being incubated with the respective PLA-PLUS and MINUS probes provided in the kit as per manufacturer’s instructions followed by the ligation and amplification steps. After the final washes, the coverslips were mounted onto slides using the Duolink® In Situ Mounting Media with DAPI (DUO82040, Sigma Aldrich). The slides were stored in the dark for at least 15 mins before being imaged using a 20x objective on a Leica Stellaris laser scanning confocal microscope (Leica GmbH, Mannheim Germany) on a DMI8 base using the LAS X software.

### Protein purification and labelling

To purify human IRSp53 or VASP from *E. coli*, the respective codon-optimized sequences were cloned into a pET28b vector, which features a thrombin-cleavable N-terminal 6xHis-tag configuration. *Escherichia coli* strain BL21 (DE3) cells was transformed with these expression vectors and grown at 37°C in Luria Bertani (LB) broth until OD_600_ of 0.5 was reached. Addition of 0.1mM IPTG was used to induce protein expression, and cells were harvested by centrifugation after 16h of expression. The cells were resuspended in 40 mM Tris-HCl, 10 mM NaP, 400 mM NaCl, 10% v/v glycerol, pH 8.0 (lysis buffer) with DNase I (Invitrogen) and cOmplete™ protease inhibitor cocktail (Roche) and lysed by sonication at 4°C. The lysate was cleared by centrifugation, and the supernatant was loaded onto two 5 ml HisTrap columns (Cytiva), pre-equilibrated in the lysis buffer. The columns were washed with 40 ml lysis buffer containing 30 mM imidazole (#I202, Sigma Aldrich). Proteins were eluted in lysis buffer supplemented with 0.5 M imidazole. Eluted proteins were treated with ULP1 SUMO protease (Thermo Fisher Scientific), after which the samples were dialyzed overnight into 40 mM Tris-HCl pH 8.0, 300 mM NaCl. The cleaved proteins were passed over the HisTrap column to remove the His-SUMO tag. The eluted proteins were collected and mixed with 2 volumetric parts of 50 mM NaP pH 6.0, 50 mM NaCl and loaded onto a HiTrap SP HP column (Cytiva). The proteins were eluted with a gradient from 9 to 55% (IRSp53) or 4 to 45% (VASP) ÄKTA pure buffer B (1 M NaCl, 50 mM Tris, pH 6.0). The purified proteins were dialyzed against 25 mM Tris-HCl pH 7.5, 300 mM NaCl, concentrated and stored at −80°C. To purify the mouse capping protein from *E. coli*, BL21 (DE3) cells were transformed with the pRSF Duet 1 (MmCPα1β2) vector (a gift from Brad Nolen, University of Oregon)^111^. The bacteria were transformed with this expression vectors and grown in LB medium at 37°C until OD_600_ of 0.5 was reached. Addition of 0.5 mM IPTG was used to induce protein expression, and cells were harvested by centrifugation after 16h of expression at 16°C. The cells were pelleted, washed with cold 1x PBS, and resuspended in extraction buffer (50 mM NaH_2_PO_4_ pH 8.0, 500 mM NaCl, 10% v/v glycerol, 10 mM imidazole, 10 mM β-mercaptoethanol, 0.5 mM PMSF and 2 protease inhibitor tablets (cOmplete, Roche)). The cells were lysed by sonication, and the lysate was clarified at 25,000xg for 30 min at 4°C. The supernatant was loaded onto a Talon metal affinity column (Clontech) and the capping protein was eluted in extraction buffer supplemented with 250 mM imidazole. Peak fractions were pooled and dialyzed overnight at 4°C against dialysis buffer (20 mM Tris pH 8.0, 50 mM NaCl, 5% v/w glycerol, 0.01% w/v NaN_3_, 1 mM DTT) using a 10 kDa molecular weight cutoff membrane dialysis tube (Spectrum Labs). The protein concentration was determined using a NanoDrop spectrophotometer, and samples were concentrated using a 10 kDa Centricon filter (Millipore). The purified capping protein was aliquoted, flash frozen and stored in the dialysis buffer at −80°C. Profilin-1(#PR02-B), fascin-1(#CS-FSC01) and Actin:rabbit skeletal muscle (#AKL95-B) were purchased form Cytoskeleton Inc. and resuspended and stored as per manufacturer’s directions.

### IRSp53 Labeling

Purified IRSp53 was dialyzed against 20mM Tris-HCl, pH 7.5, 150 mM NaCl. 300 µM IRSp53 was treated with a 100x excess of Tris(2-carboxyethyl)phosphine (TCEP) and a 10x molar excess of C5-Maleimide-AF488 (#A10254, Thermo Scientific). A NAP-5 column (GE healthcare) was used to separate the labeled protein from the free dye. Individual fractions were collected and absorbance at 488 nm and 280 nm was used to determine the concentrations of the dye and the protein. These values were used to calculate labeling efficiency.

### Polyphosphate end-labelling

PolyP-300 (300-mer chain) was labelled with a custom synthesized peptide, KASASHHHHHH (Genscript) following the protocol outlined by ^50^. In short, 50 µM polyP-300 chains (equivalent to 100 µM polyP ends) in 100 mM MOPS-NaOH, pH 8.0 was incubated with 8 mM of the custom peptide in the presence of 150 mM 1-Ethyl-3-(3-dimethylaminopropyl) carbodiimide hydrochloride (EDAC) for 1 h at 37°C. Excess peptide and EDAC were removed by using a 5mL Zeba^TM^ Spin desalting column using the same buffer. His-tagged polyP-300 was bound to HisPur-NiNTA resin beads (#88221, Thermo Scientific) packed in a column and pre-equilibrated with 80 mM MOPS-NaOH, pH 8.0, 300 mM NaCl, 10 mM imidazole. The column was washed with four column volumes (CV) of pre-equilibration buffer. His-polyP was eluted using 80 mM MOPS-NaOH, pH 8.0, 300 mM NaCl, 500 mM imidazole. The eluant was pooled and buffer exchanged into 100 mM MOPS-NaOH buffer, pH 8.0, 300 mM NaCl, 3 mM imidazole using a 5 mL Zeba^TM^ Spin desalting column after which it was loaded onto a second column packed with HisPur-NiNTA resin beads (#88221, Thermo Scientific) pre-equilibrated as before. The column was washed and His-tagged polyP was eluted as done earlier. The resulting eluant was buffer exchanged into 20 mM HEPES, pH 8.0 as before, and concentrated using an Amicon 3K Ultra-0.5ml centrifugal filter (EMD Millipore). The polyP concentration was measured using the DAPI-based fluorescence assay (see above).

### Protein turbidity assays, *in vitro* condensate formation and FRAP measurements

10 µM purified IRSp53 in 50 mM KPi, pH 7.4, 100 mM KCl was incubated in the absence or presence of increasing concentrations of various length polyP-chains. Turbidity measurements were conducted in clear-walled 96-well plates. Turbidity was measured at 360 nm using a Tecan M1000 (Tecan). Individual samples were imaged in 16-well CultureWell slides (Grace Bio-Labs, #GBL112358) treated with 5% w/v pluronic acid F-127 (Sigma, P2443) overnight and washed with ddH_2_O prior to sample preparation. For the phase diagram, purified IRSp53 was mixed with polyP-300 at the indicated concentrations in 50 mM KPi, pH 7.4, 100 mM KCl. The samples were prepared in 16-well CultureWell slides (Grace Bio-Labs, #GBL112358), which had been treated overnight with 5% w/v pluronic F-127 (Sigma, P2443) and washed extensively with ddH2O. Imaging was performed on a Leica SP8 confocal microscope using the brightfield channel. To visualize co-condensate formation, 100 µM polyP-300 spiked with 2% labelled polyP300-647 and 10 µM purified IRSp53 spiked with 2% labelled IRSp53-488 were incubated in 50 mM KPi, pH 7.4, 100 mM KCl. The samples were prepared as before in pluronic acid-treated chambered. CultureWell slides. Samples were imaged on the ECHO Revolution (ECHO) using the 20x objective. FRAP measurements of the droplets were conducted on a Leica SP8 inverted microscope with 100x oil immersion objective, driven by LAS X software (Leica GmbH, Mannheim, Germany). The Zoom-In mode was used. Partial fluorescence bleaching of similarly sized droplets was achieved by using the 488 nm laser at 100% intensity. Two pre-bleach images were acquired, followed by photobleaching and fluorescence recovery measurements every 5 sec for 100 sec. The fluorescence intensity was measured by LAS X software. Recovery data were plotted and fitted against a one-phase association curve with GraphPad Prism.

### Giant unilamellar vesicle (GUV) preparation

GUVs were prepared using a modified phase transfer protocol optimized for high encapsulation efficiency and membrane uniformity^112^. Lipid films were generated by dissolving a lipid mixture of 50.0 mol% porcine brain polar lipid extract (Avanti), 0.1 mol% rhodamine B-labeled phosphoethanolamine (Rhod-PE) (Avanti) and 44.9 mol% 1-palmitoyl-2-oleoyl-sn-glycero-3-phosphocholine (POPC) (Avanti) together with either 5.0 mol% 18:1 1,2-dioleoyl-sn-glycero-3-[(N-(5-amino-1-carboxypentyl)-iminodiacetic acid)succinyl] (DGS-NTA (Ni)) (Avanti), L-α-phosphatidylinositol-4,5-bisphosphate (Brain, Porcine) (PIP2) (Avanti) or 18:1 1,2-dioleoyl-sn-glycero-3-phospho-L-serine (DOPS) (Avanti) in chloroform to a final concentration of 0.44 mM. The lipid solution was deposited in a clean glass vial and evaporated under a gentle argon stream while rotating the vial to ensure uniform film distribution. Residual solvent was removed by placing the vial under vacuum for ≥1 hour. The dried lipid film was resuspended in 1.1 ml of mineral oil (Sigma) by vortexing for 1 min, followed by ultrasonication at 55°C for 20 min in an ultrasonic cleaner (Vevor) to ensure complete lipid dispersion. To form a stable oil–water interface, 100 µl of outer aqueous solution (310 mM glucose) was added to a 1.5 ml microcentrifuge tube. Next, 100 µl of the lipid-oil solution was carefully layered on top using a slow pipetting motion against the inner wall to minimize turbulence. The interface was allowed to equilibrate for 1 hour at RT. Separately, 20 µl of inner aqueous solution (310 mM glucose) containing 10% v/v Opti-prep (Sigma-Aldrich), was added to a fresh tube. This solution was gently overlaid with 200 µl of the lipid–oil mixture, and the two phases were emulsified by slow pipetting until the emulsion became uniformly colored. The emulsion was then slowly added onto the preformed oil-water interface. The assembled gradient was centrifuged at 2,500xg for 10 min at RT using an Eppendorf 5415R centrifuge rotor. Following centrifugation, the upper oil phase and aggregated material were carefully aspirated using a 200 µl pipette. Once the remaining oil volume reached ∼50 µl, finer removal was performed using a 10 µl pipette tip in a slow circular motion, changing tips to avoid contamination. The GUV pellet at the bottom of the tube was resuspended by gentle pipetting (10–15 times) until the solution appeared homogeneously colored.

### Preparation of polyP-enriched GUVS and assaying protein recruitment

Ni-NTA containing GUVs and, as negative controls, control and PIP2-enriched GUVs were incubated with 2.5 µM His-tagged polyP for 1 hour at RT. The GUVs were pelleted by centrifugation at 100xg for 10 min. GUVs were incubated with 10 µg/ml of PPXBD-GFP for 1hr at RT to quantify polyP binding on the membrane. To determine the recruitment of IRSp53, the various GUVs were incubated with 50 nM IRSp53-AF488 for 1 hr at RT in a black walled 96-well plate with a coverslip bottom (Ibidi). The GUVs were imaged on a Leica SP8 confocal microscope and resulting AF488 fluorescence on the membrane was quantified on FIJI to determine the extent of IRSp53 recruitment. Protrusion formation assays were conducted following the protocol described in^42^. Briefly, the 30 µl of GUVs were incubated with 16 nM IRSp53, 4 nM VASP, 250 nM fascin, 600 nM profilin, 25 nM capping protein, 500 nM monomeric actin and 660 nM Phalloidin-647 in 1xF-buffer (10 mM Tris HCl, pH 7.5, 50 mM KCl, 2 mM MgCl_2_) supplemented freshly with 1 mM ATP (#R0441, Thermo Scientific). The GUV-protein mixture was incubated at RT for at least 30 min and imaged on a Leica SP8 confocal microscope as before. To determine the fraction of GUVs containing inward protrusions, the number of GUVs with at least a single membrane protrusion was counted manually and divided by the total number of GUVs in the same field of view. All images were blinded prior to analysis.

### Trans-well migration and invasion assays

The trans-well *migration assay* was conducted with the CytoSelect™ 24-Well Cell Migration Assay, 8 μm (#CBA-100, CellBio Labs) using the manufacturer’s protocol. The trans-well *invasion assay* was conducted using the CytoSelectTM Cell Invasion Assay (#CBA-110, CellBio Labs) following the manufacturer’s protocol. In each case, ∼200,000 cells in serum-free media were seeded into the respective trans-well chambers. Media with serum was added to the outside of the chamber and cells were allowed to migrate or invade for 4 hours. After that time, all cells that remained in the chamber were removed. Cells that were able to migrate through the membrane were fixed and stained with 0.1% v/v crystal violet (Sigma Aldrich). The membrane was imaged and cells were counted using the Cell Counter plugin on FIJI.

### Cell adherence assays

To measure cell adherence, ∼10,000 cells per well were plated in a 96-well culture plate and allowed to adhere for 15 mins, 30 mins, 1 hr, 1.5 hrs, 2 hrs, 3 hrs or 4 hrs. After the respective times, the media was removed and the wells were washed thoroughly with pre-warmed media to remove all non-adherent cells. The remaining cells were allowed to further adhere overnight after which the cells were fixed and stained with 0.1% w/v crystal violet. After washing with ddH2O to remove all excess crystal violet from the wells, the cells were lysed and the absorbance was measured at 562 nm using a Tecan M1000 plate reader. In one well per condition, all cells were allowed to adhere. This value was set to a 100% for that condition. A well containing media alone was treated was used to set the baseline for the absorbance measurements.

### Lipid Nanoparticle (LNP) loading and delivery to fibroblasts

100 µl BODIPY labelled loadable lipid nanoparticles (LipidLaunch™ BODIPY SM-102 LNP; Cayman Chemical) were incubated with 20 µl of either water or 10 mM or 50 mM polyP-300 (kindly provided by T. Shiba, RegeneTiss, Japan) in water. Upon mixing the samples, 40 µl of encapsulation buffer provided in the kit was added and the mixture was incubated at RT for 10-15 minutes. This mixture was divided amongst three wells of a six-well plate seeded with ∼150,000 cells each in a total volume of 2 ml media. After 24 hours incubation at 37°C, the uptake of LNPs was validated by visualizing BODIPY puncta inside the cells using the ECHO Revolution at 10x magnification.

### Tumor mice models

Mouse husbandry and procedures were in accordance with protocols approved by The University of Michigan Animal Care and Use Committee (IACUC protocol: PRO00012211). Wildtype FVB/NJ (JAX #001800), MMTV-PyMT (JAX#002374), and C3(1)-Tag (JAX#013591) mouse strains were obtained from the Jackson Laboratory and maintained in the FVB/n background. Statistical tests were not employed to determine sample size before experiments as the effect of treatment was unknown. There was a random assignment of littermates to experimental groups.

### Organoid isolation and 3D culture

Primary organoids were isolated by following previously published protocols^64^. In short, either healthy mammary tissue or tumors were dissected out of the respective mice once the tumor reached ∼1.5 cm in width and finely minced 50-100 times with a scalpel. The minced tissue was incubated in a collagenase solution (DMEM-F12 (#10-565-018, Fisher Scientific), 5% v/v FBS (#F4135, Sigma-Aldrich), 50 μg/mL gentamycin(#15750060, Thermo Scientific), 5 μg/mL insulin (#I9278, Sigma-Aldrich), 2 mg/mL collagenase (#C2139, Sigma Aldrich) and 2 mg/mL trypsin (#VWRV0458, VWR)) for 1h at 37°C with shaking. The digested tissue was centrifuged at 1,300 rpm for 10 mins after which the pellet was resuspended in a DNase solution (2 U/ml) in DMEM-F12 and incubated at RT for 5 mins. After quenching DNase activity by addition of excess media, the solution was centrifuged at 1,300 rpm for 10 mins. The pellet was resuspended in DMEM-F12 and larger, undigested tissue was allowed to pellet via gravity after which the supernatant was transferred to a new tube. The organoids were separated from the immune and stromal cells by a series of differential centrifugations at 1,300 rpm for 10 sec each. The resulting organoids were used for further assays. To deliver polyP via LNPs, 20 µl of buffer or polyP-loaded LNPs were added per 1,000 organoids and incubated in a 24-well plate coated with anti-adherence rinsing solution (#07010, STEMCELL technologies) for 24 hours at 37°C in 5% CO_2_. The treated organoids were spun down, washed with DMEM-F12 and embedded in 3D fibrillar rat tail collagen I matrix (#354236, Corning) at a density of 1-2 organoids per μl and plated in a 24-well coverslip bottom plate (#82426, Ibidi). The collagen gels were allowed to polymerize for 0.5–1 h at 37°C, after which organoid growth media (DMEM-F12, 1% w/v penicillin–streptomycin (#SV30010, Cytiva), 1% w/v insulin–transferrin–selenium–ethanolamine (51-500-056, Fisher Scientific)) was added. After 24 hours, half of the organoid-embedded gels were collected for cryo-sectioning and RNA isolation (Day 0 organoids) (see below). The remaining organoid-embedded gels were treated with 2.5 nM basic fibroblast growth factor (FGF2) (#F0291, Sigma-Aldrich) and maintained for 7 days (Day 7 organoids) to measure invasion (see below).

### Cryosectioning and immunofluorescence staining of organoids

Collagen gels with embedded organoids were fixed in 4% v/v PFA (#1578100, Electron Microscopy Sciences) overnight at 4°C. Gels were washed 3-times with PBS before the gels were embedded in optical cutting temperature compound (OCT) (#4583; Tissue-TEK) and frozen in a slurry of dry-ice and ethanol. The OCT blocks were stored at – 80°C. The gels were sectioned at 50 μm thickness onto Superfrost Plus Gold Microscope slides at −20°C using a cryostat (Leica). The OCT compound was washed away from the slides with three PBS washes of 5 min each. The gel slices were then permeabilized for 1 h with 0.5% v/v Triton X-100 (T8787, Sigma-Aldrich) in PBS and blocked in PBS supplemented with 10% v/v FBS (F4135, Sigma-Aldrich) and 1% w/v BSA (A3059, Sigma-Aldrich) at RT for 2 h. The slices were incubated in a mix of mouse anti-E-cadherin (14472S, Cell Signaling Technologies) at the recommended dilution and 10 µg/ml PPXBD-mCherry/GFP in PBS buffer containing 1% v/v FBS, 1% w/v BSA, 0.1% v/v Triton X-100 and incubated overnight at 4°C. After three PBS washes, the slides were incubated with goat anti-mouse IgG (H+L) secondary antibody, Alexa Fluor 568 (#A11004, Fisher Scientific) for 2 h at RT. The slides were washed and stained with 1 μg/ml DAPI (#D1306, Thermo Fisher Scientific) for 15 mins to stain the nucleus. Afterwards, the slides were washed again with PBS and mounted with Prolong Gold mounting media (9071S, Cell Signaling Technology) and covered with 12 mm circular coverslips (72230-01, Electron Microscopy Sciences). Mounted coverslips were sealed with nail polish and imaged after at least 24 hours post-mounting. Mounted cells were visualized using a 63x oil objective on a Leica Stellaris laser scanning confocal microscope (Leica GmbH, Mannheim Germany) on a DMI8 base using the LAS X software.

### 3D-Organotypic invasion measurements

Organoids embedded in collagen were allowed to invade for 7 days after which they were fixed in 2% v/v PFA for 20 mins at RT. The fixed gels were washed with 1xPBS and incubated with 1.32 µM Phalloidin-647 (#8940, Cell Signaling Technology) and 1 μg/ml DAPI (#D1306, Thermo Fisher Scientific) for 30 mins. The gels were washed 3-times with PBS and stored in PBS until imaging. Whole organoids were visualized using a 4x objective on a Leica Stellaris laser scanning confocal microscope (Leica GmbH, Mannheim Germany) on a DMI8 base using the LAS X software. Invasiveness was represented as inverse circularity measurements of maximum intensity projections of the actin channel generated by the FIJI software.

### Organoid RNA-seq and analysis

About 1,500 organoids per mouse, condition and timepoint were embedded at a density of ∼1.5 organoids/µl of collagen. RNA was extracted from Day 0 and Day 7 organoids isolated from PyMT-MMTV mice; n=4 mice per treatment. Briefly, the collagen was dissolved by incubating the gels in 5 ml of a 2 mg/ml collagenase-PBS solution at 37°C for 10 mins. The released organoids were pelleted and washed with PBS. RNA was extracted using the Invitrogen Trizol plus RNA purification kit (Thermo: 12183555). Purified RNA was sent to Plasmidosaurus for poly(A) selection library prep and sequencing. Raw sequencing files were subjected to Fastqc to assess data quality using default parameters ^113^. Illumina adaptors were removed using Trimmomatic ^114^ by performing an initial ILLUMINACLIP step and an average read quality of 20. Sequences were aligned to the mm39 (GRCm39) mouse genome and gene abundance was quantified with Salmon^115^ on single-end read mode and achieved a mapping rate of 77% or higher for all samples. Gene counts were imported into RStudio with TXIMport ^116^. DESeq2_1.52.0 ^117^ was used to compare gene expression profiles between conditions with a multifactor design to account for all experimental groups and experimental batches. Significant differentially expressed genes were defined using a p-value cutoff of p<0.05 and a Log_2_FC>1.5 or Log_2_FC<-1.5. For gene set enrichment analyses, fgsea was used with MSigDB hallmark gene sets and default settings. The code is available upon request.

