## Supplemental Information for "The filopodial scaffold polyphosphate dictates cell adhesion-*versus*-invasion decisions"

Supplementary Figure 1

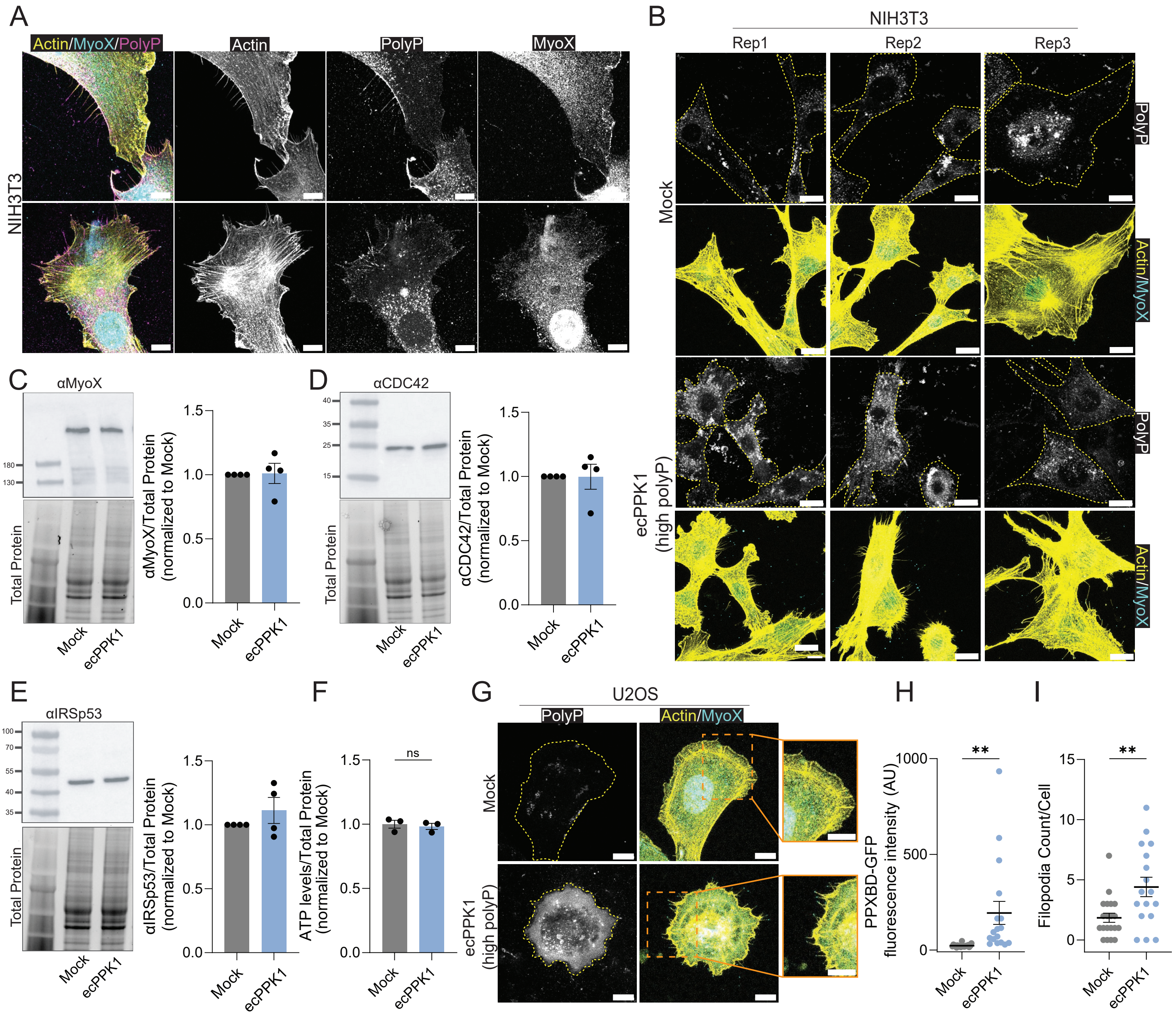


**Figure S1. Increase in polyP levels increases filopodial length and dynamic properties (related to Figure 1). (A)** NIH3T3 mouse embryonic fibroblasts stained for polyP, actin and MyoX as shown in Fig. 1A. **(B)** Cells from three different passages of mock- or EcPPK1-expressing NIH3T3 fibroblasts were stained for actin (yellow) and MyoX (cyan) and used for the quantification of filopodia number and length shown in Fig. 1F, G. **(C-E)** Lysates from mock- or EcPPK1-expressing NIH3T3 fibroblasts were analyzed by Western Blot, using antibodies against **(C)** MyoX, **(D)** CDC42 or **(E)** IRSp53. Densitometry plots were used to quantify band intensities, which were normalized against the total protein signal according to the stain-free PAGE (lower panels). n=4 biological replicates. **(F)** ATP levels in mock- or EcPPK-transfected NIH3T3 fibroblasts after 24 hours of expression normalized to total protein. Each data point represents the average ATP levels from ~15,000 cells normalized to total protein signal obtained from running lysates on na stain free gel. n=3 biological replicates. **(G)** Mock- or EcPPK1-transfected U2OS osteosarcoma cells were fixed and stained for polyP (gray), actin (yellow) or MyoX (cyan). Quantification of the **(H)** fluorescent polyP signal or (I) filopodia number of cells shown in (E). Each datapoints in panel H represents polyP intensities of individual cells as quantified by measuring PPXBD-GFP fluorescence intensity. Mean fluorescence intensity of mock transfected cells was set to 1. **(I)** Each data point in panel I represents the average MyoX-positive filopodia count of 5-25 cells. Unpaired t-test was used to determine statistical significance between samples. * $p < 0.05$, *** $p < 0.001$, **** $p < 0.0001$.

Supplementary Figure 2

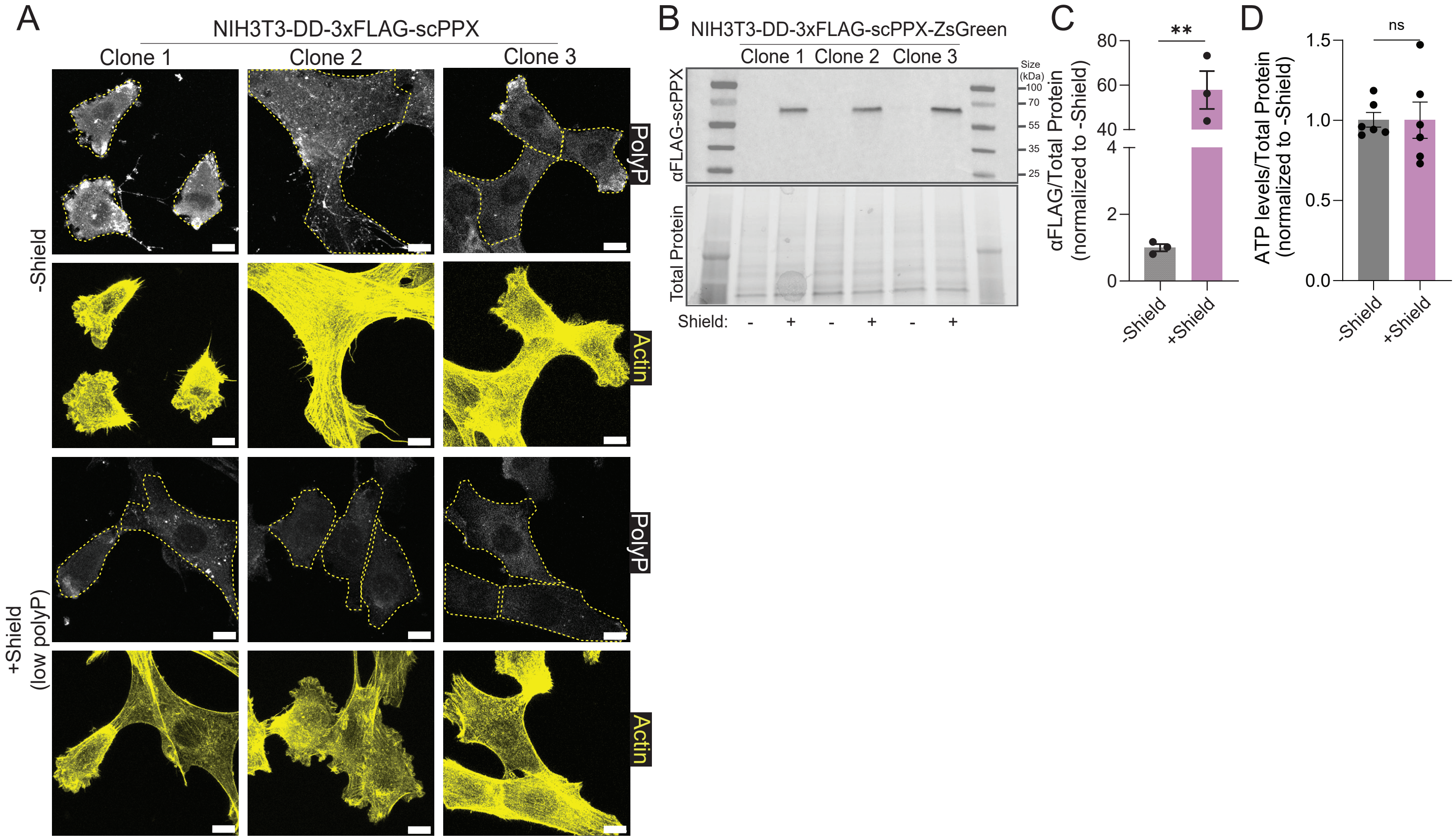


**Figure S2. Decrease in cellular polyP levels increases filopodia dynamics (related to Figure 2).** **(A)** Three different clonal populations of stably transfected NIH3T3-DD-3xFLAG-ScPPX cells were fixed and stained for polyP (gray) or actin (yellow) in the absence or presence of the stabilizer Shield. These images were used for quantification of filopodia length and number shown in Fig. 2. **(B)** Expression of 3xFLAG-ScPPX in three clonal cell lines was tested after 24h of incubation in the presence of Shield. Western blot analysis of lysates using anti-FLAG tag antibodies is shown in upper panel. Coomassie-stained SDS-PAGE of the respective samples is shown in lower panel. **(C)** Quantification of the band intensities shown in (B, upper panel) and normalized against the total protein shown in (B, lower panel). **(D)** ATP levels in NIH3T3-DD-3xFLAG-ScPPX fibroblasts after 24 hours of incubation in the absence or presence of Shield normalized to total protein. Each data point represents the average ATP levels from ~15,000 cells normalized to total protein signal obtained from running the respective lysates on a stain-free gel; n=3 biological replicates. Scale bars for main images 10 µm. Unpaired t-test was used to determine statistical significance between samples. ** $p < 0.01$; ns, not significant.

### Supplementary Figure 3

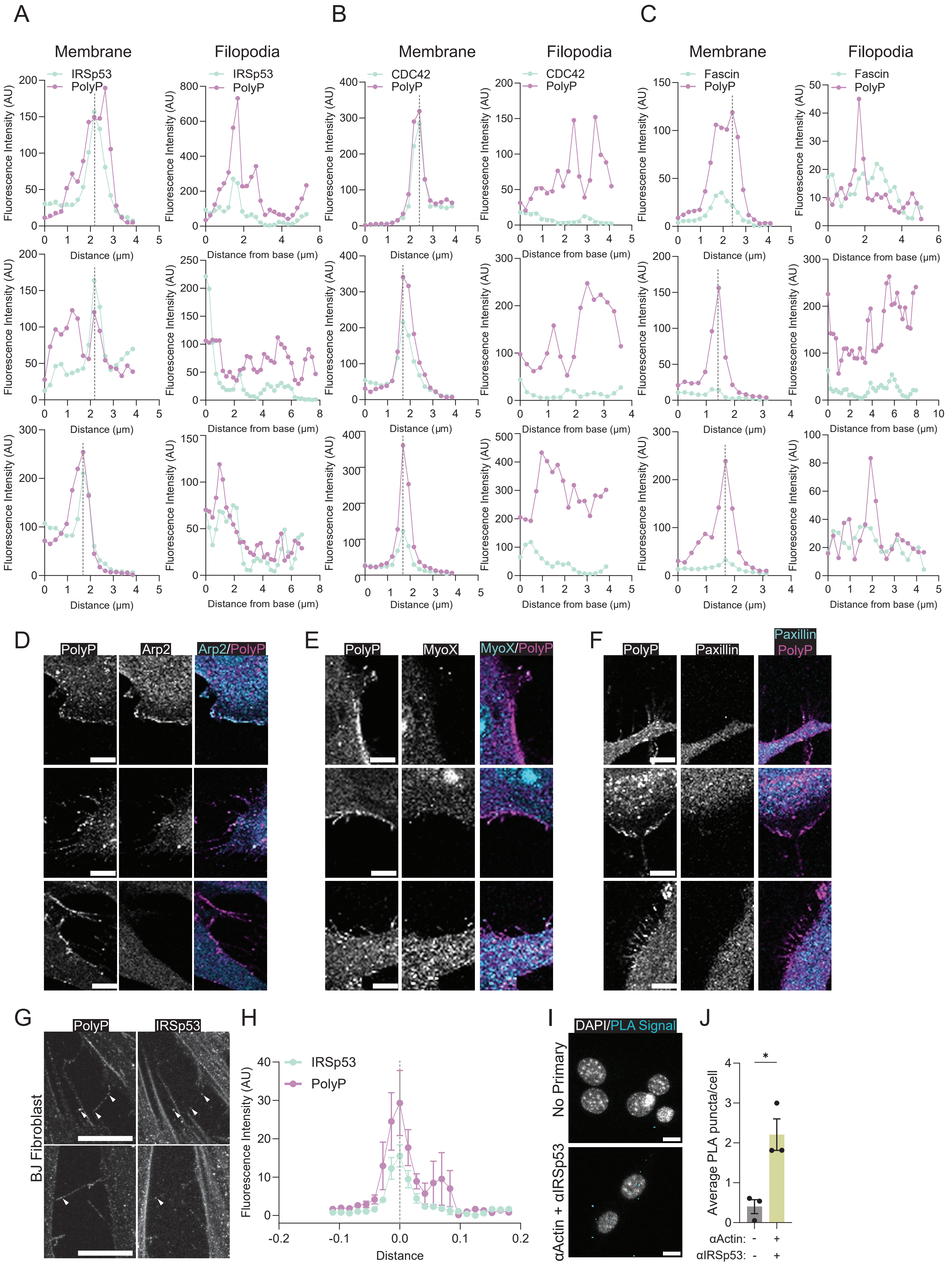

**Figure S3. Figure 3. PolyP is a hitherto unknown component of mammalian filopodia (related to Figure 3).**

**(A-C)** Three different preparations of NIH3T3 fibroblasts were fixed and stained for polyP (PPXBD-GFP) and either IRSp53, CDC42 or fascin. Representative intensity profiles from each replicate were analyzed for colocalization of (A) IRSp53, (B) CDC42, (C) fascin on the membrane or within the filopodium. (D) Co-staining of fixed NIH3T3 cells with PPXBD-GFP and either **(D)** Arp2, **(E)** MyoX, or **(F)** paxillin. **(G)** BJ fibroblasts were fixed and co-stained for polyP (PPXBD-GFP) and IRSp53. **(H)** Quantification of the intensity profiles of individual polyP puncta and the corresponding IRSp53 intensity in 10 different filopodia shown in (G). A maximum of two puncta per cell were quantified. **(I)** PLA of fixed NIH3T3 fibroblasts. Cells were incubated with either buffer (no primary) or mouse anti-actin and rabbit anti-IRSP53 before incubation with oligonucleotide linked secondary antibodies. The PLA amplification signal is depicted as cyan puncta. **(J)** Quantification of PLA signals shown in (I). Each data point represents the total number of PLA puncta in a field of view divided by the total number of cells in that same field of view. Scale bars for all images 10 µm. Unpaired t-tests were used to determine statistical significance between samples. * $p < 0.05$, *** $p < 0.001$, **** $p < 0.0001$. Quantitative image analysis was conducted blinded.

Supplementary Figure 4

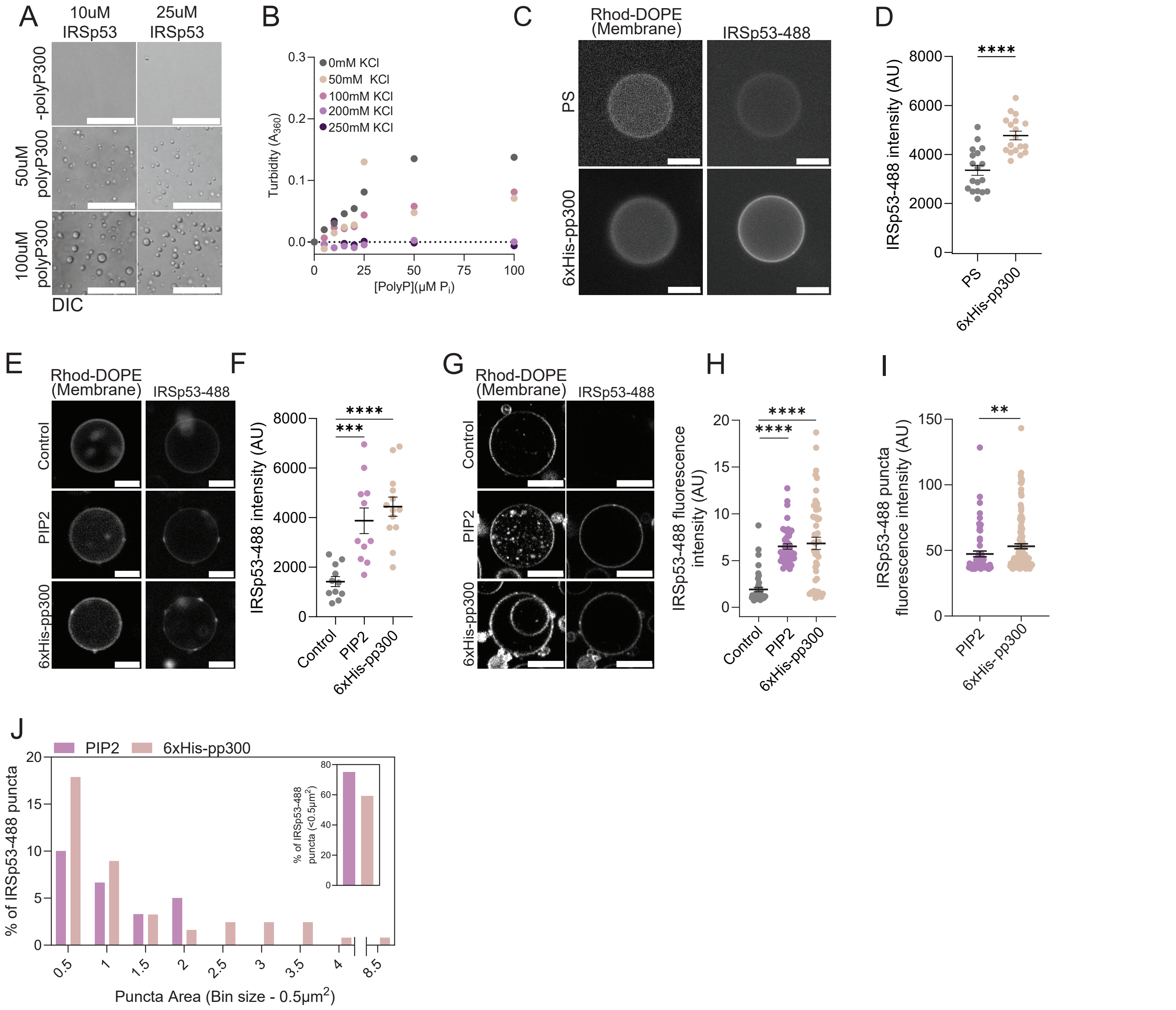


**Figure S4. Membrane-bound polyP recruits IRSp53 and supports ex vivo filopodia formation (related to Fig. 4)**
**(A)** DIC images of solutions containing 10 or 25 μM IRSp53 in 50 mM KPi, pH 7.4, 100 mM KCl in the absence or presence of the indicated concentrations of polyP-300 (concentration in Pi-units). Scale bars: 100 μm. **(B)** Salt sensitivity of IRSp53-polyP condensates. Condensates were formed by incubating 10 μM IRSp53 in 50 mM KPi, pH of 7.4 with increasing concentrations of polyP-300 (in Pi-units) in the absence or presence of increasing amounts of KCl. The turbidity of the solution was measured at $OD_{360}$. (C) IRSp53-488 signal of control or PS-enriched GUVs after 15 min incubation with 50 nM IRSp53-488. The membranes were visualized using the rhodamine-DOPE signal. Scale bars: 10 μm (D) Quantification of the IRSp53 signal on the membranes of GUVs shown in **(C)**. **(E, F and G, H)** Replicates of experiments shown in Fig. 4I,J using two different GUV preparations. **(I)** Signal of IRSp53-488 puncta on the membrane of PIP2- enriched or polyP-enriched GUVs after 15 min incubation with 50 nM IRSp53-488 using data from all three replicates for input. Each data point represents the mean fluorescence intensity of a single IRSp53-488 cluster on the surface of a GUV; n= >60 clusters per condition. **(J)** Size distribution of IRSp53-AF488 clusters on polyP-enriched versus PIP2-enriched membranes using a 0.5 μm bin size. Inset: Percent of IRSp53-AF488 puncta with a size of < 0.5 $\mu m^2$. Scale bars in panels C, E, G:10 μm. One way ANOVA analysis was used to determine statistical significance between samples. * $p < 0.05$, ** $p<0.01$, *** $p < 0.001$, **** $p < 0.0001$. Quantitative image analysis was conducted blinded.

Supplementary Figure 5

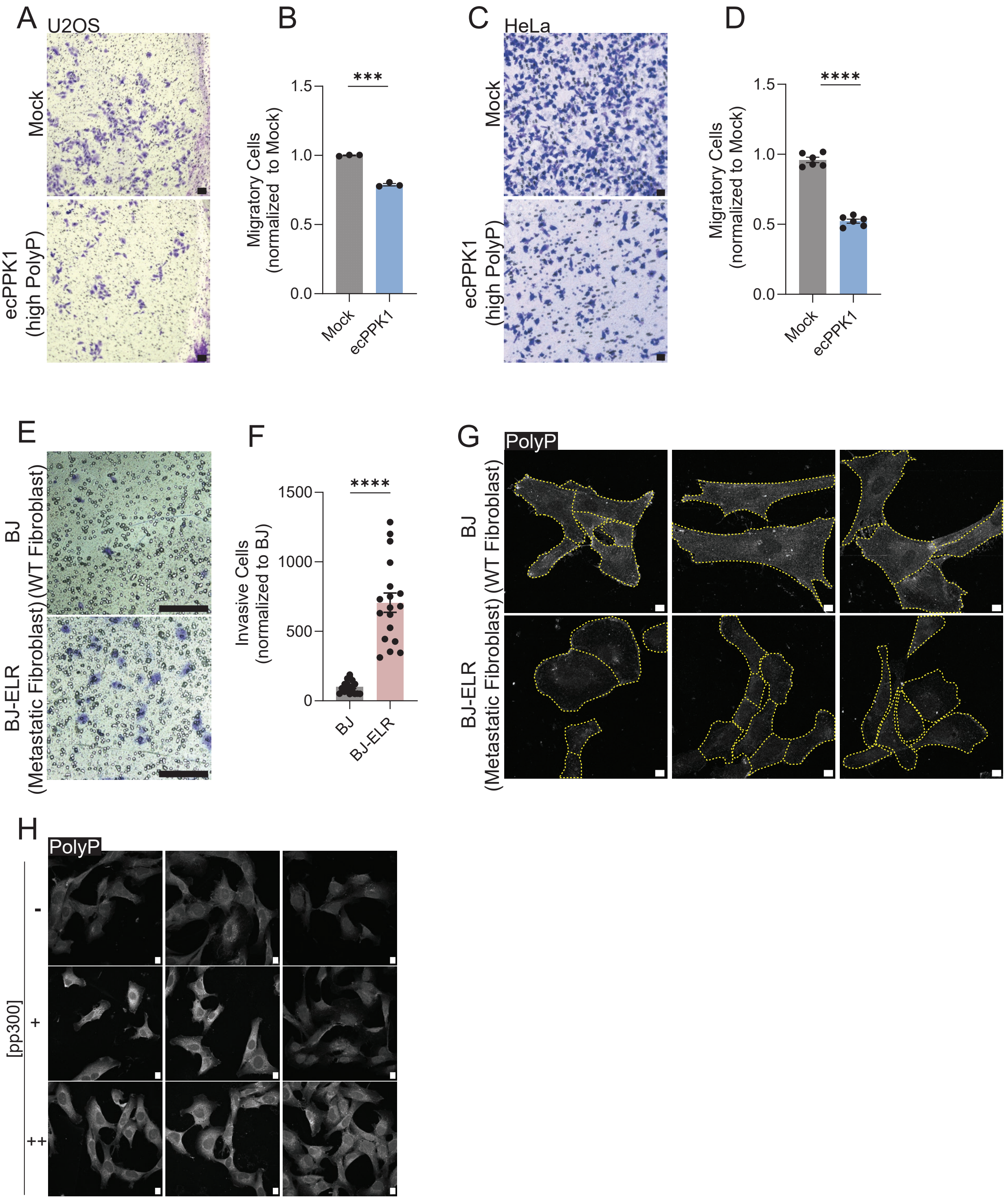


**Figure S5. PolyP promotes cell adhesion and reduces trans-well migration (related to Figure 5) (A, C)** Trans-well migration assay of mock- or EcPPK1 transfected (A) U2OS human osteosarcoma cells or (C) HeLa cervical cancer cells. For details see Fig. 5A. **(B,D)** Quantification of data shown in (A,C). **(E)** Trans-well invasion assay of BJ and BJ-ELR fibroblasts. For details see Fig. 5O. **(F)** Quantification of data shown in **(E)**. Scale bars: 100 µm for boyden chamber assays – 100um. Unpaired t-tests were used to compare pairs of samples whereas one way ANOVA was used to analyze samples with multiple comparisons with a p-value threshold of 0.05. **(G)** Biological replicates of Fig. 5G. **(H)** Biological replicates of Fig. 5J.

Supplementary Figure 6

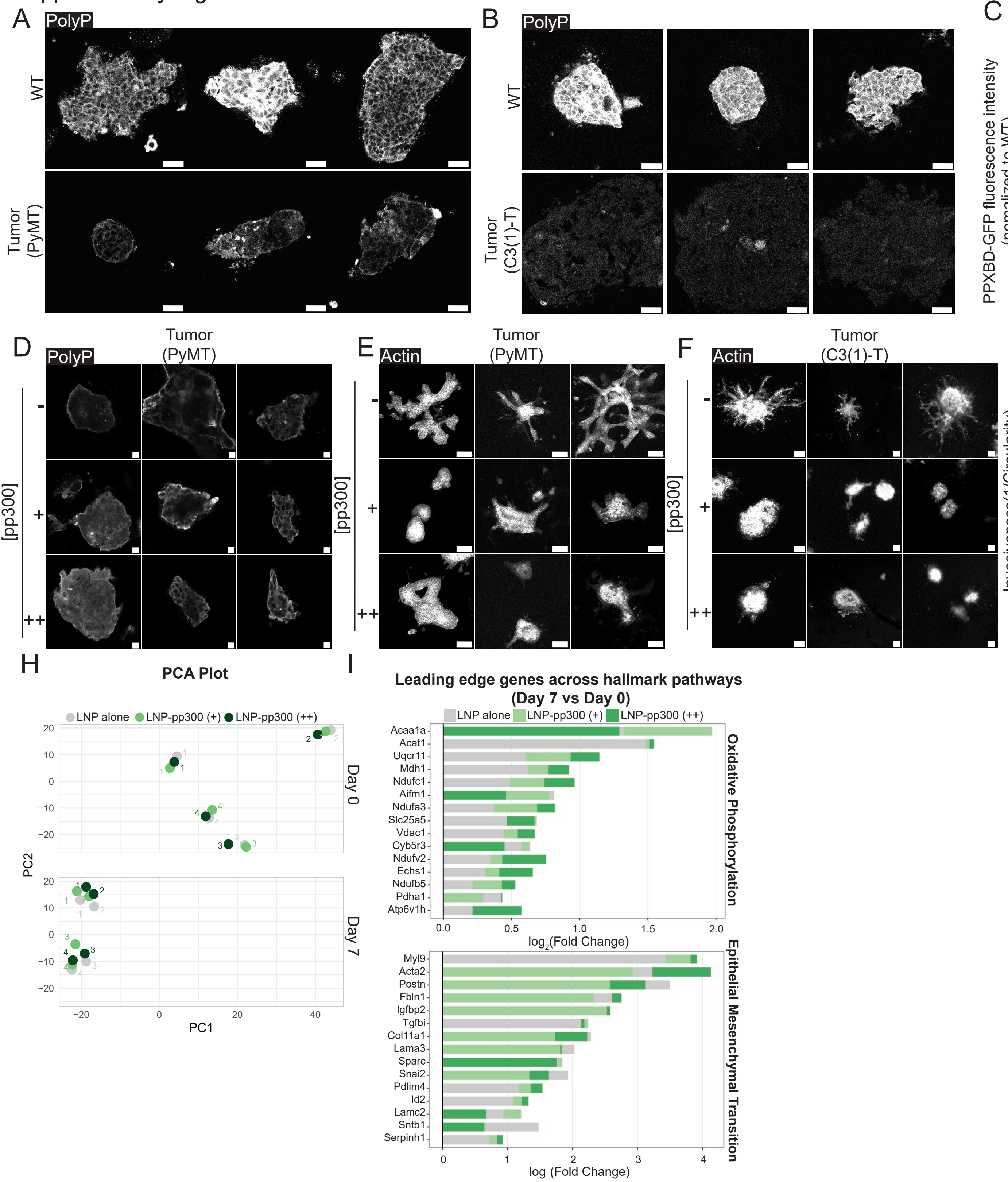

**Figure S6. Restoration of polyP levels in organoids reverses metastatic features (related to Figure 6)**
**(A)** Additional fluorescence images of organoids derived from healthy mammary tissue or MMTV-PyMT tumor tissue using three different mice each; cryosections were stained for polyP using PPXBD-GFP. See Figure 6A for more details. **(B)** Fluorescent images of healthy mammary tissue-derived or C3(1)-T tumor-derived organoids one day after seeding in collagen I gels. Staining with PPXBD-GFP was used to visualize polyP (gray). n=3 mice per condition. **(C)** Quantification of the immunofluorescence images shown in (B). Each data point represents the average PPXBD-GFP fluorescence intensity of a single organoid **(D)** Additional fluorescence images of MMTV-PyMT derived organoids pre-treated with buffer- or polyP-300 loaded LNPS; the cryosections were stained for polyP using PPXBD-mCherry. See Figure 6E for more details. **(E)** Additional images from 3D-invasion assay of MMTV-PyMT derived organoids pre-treated with buffer- or polyP-300 loaded LNPS. See Figure 6G for more details. **(F)** 3D-invasiveness of C3(1)-T tumor-derived organoids pre-treated for 24h with buffer-loaded (-) or polyP-loaded (+, 10 mM polyP-300; ++, 50 mM polyP-300) LNPs before seeding in collagen I gels. After 7 days in FGF-supplemented media, the organoids were fixed and stained for DAPI and actin. n=3 mice. **(G)** Inverse circularity (IC) as quantitative readout for invasiveness of organoids shown in (F). Each data point represents IC of a single organoid. Average IC of organoids treated with buffer loaded (-) LNPs organoids is set to 1; n=3 mice. **(H)** Principal Component Analysis (PCA) plot of RNA-Seq data. **(I)** Stacked bar plot representing the log2(FC) of select genes belonging to the leading edge of the indicated hallmark pathways between at D7 vs D0 of MMTV-PyMT derived organoids pretreated for 24h with buffer-loaded (-) or polyP-loaded (+, 10 mM polyP-300; ++, 50 mM polyP-300) LNPs before seeding in collagen I gels.
